# Ratio-preserving super-resolution FLIM for quantitative live-cell lifetime imaging

**DOI:** 10.64898/2026.10.25.761648

**Authors:** Chenyang Li, Linglong Li, Yiwen Cheng, Wenqing Lu, Wen Gou, Haoyu Li, Qi Zou, Dilizhatai Saimi, Zhixing Chen, Changliang Guo, Bei Liu, Xiaoshuai Huang, Junchao Fan, Liangyi Chen

## Abstract

Super-resolution (SR) fluorescence lifetime imaging promises nanoscale maps of molecular environment and function, but photon-limited acquisition and computational reconstruction can distort lifetime estimates, particularly for ratio-derived frequency-domain measurements under SR conditions. Here we introduce structured-illumination frequency-domain FLIM (SIM-fdFLIM), an SR lifetime imaging platform that combines commercial dual-tap fdFLIM detection with ratio-preserving reconstruction. SIM-fdFLIM uses direct-current (DC)-stabilized lifetime estimation to improve reconstruction robustness under low signal-to-noise conditions, followed by **<u>r</u>**atio-domain **i**ntegrated **<u>v</u>**ariational **<u>e</u>**stimation (**RIVE**) to correct channel misregistration and suppress noise amplification after SR reconstruction. The method achieves ∼100 nm spatial resolution in lifetime maps and enables live-cell imaging at 2.8 Hz over a 56 × 56 μm² field of view. Across representative fluorophore standards, RIVE reduces lifetime standard deviation by approximately threefold relative to non-RIVE SIM-fdFLIM while preserving mean lifetime agreement with calibrated widefield fdFLIM. In living cells, SIM-fdFLIM resolves mitochondrial cristae-scale lifetime heterogeneity, mitochondrion-to-mitochondrion functional dynamics, nanoscale lipid-environment variations and perturbation-sensitive organelle-contact signatures. These results establish SIM-fdFLIM as a ratio-preserving platform for quantitative SR lifetime imaging, extending live-cell FLIM into a regime that combines nanoscale spatial resolution, Hz-rate acquisition, and preserved lifetime fidelity.

---

Super-resolution (SR) microscopy has transformed biological imaging by resolving subcellular structures beyond the diffraction limit^1–5^. However, most SR modalities primarily report fluorescence intensity and therefore provide structural information rather than quantitative functional readouts. Many biological processes, including molecular interactions, metabolism, membrane organization and organelle physiology, are governed not only by molecular position but also by local chemical and physical environments^6–8^. Fluorescence lifetime imaging microscopy (FLIM) provides such functional contrast by measuring the excited-state decay of fluorophores, which is relatively insensitive to fluorophore concentration, illumination nonuniformity and detection efficiency^9,10^. Extending FLIM into the SR regime would therefore enable quantitative functional imaging at the length scale of organelle subdomains, molecular assemblies and membrane contact sites.

Several SR-FLIM strategies have been developed, including stimulated emission depletion (STED) FLIM, image-scanning FLIM, and localization-based lifetime microscopy^11–14^. These approaches have demonstrated the value of combining lifetime contrast with nanoscale spatial resolution, but they typically impose trade-offs among spatial resolution, acquisition speed, field of view, photon dose and lifetime precision^9,15^. Point-scanning implementations provide high spatial resolution but are limited in throughput and can be challenging for fast live-cell functional imaging^11^. Localization-based approaches can achieve high spatial precision but require long acquisition and are generally not suited to dense, dynamic lifetime measurements^14^. Structured illumination microscopy (SIM) offers an attractive alternative because it provides widefield acquisition, moderate illumination dose and approximately twofold resolution enhancement, making it compatible with live-cell imaging^16–18^. A recent EO-SIM-FLIM implementation has established the feasibility of SR lifetime imaging in living cells. However, because lifetime recovery in such approaches depends on preserving quantitative relationships among multiple temporally gated or modulated image channels, maintaining lifetime fidelity during high-speed, photon-limited SR reconstruction remains challenging^19^.

More broadly, many functional microscopy modalities do not directly measure a single intensity image, but derive quantitative parameter maps by combining multiple image channels, temporal gates, spectral bands or polarization states. In such cases, SR reconstruction must preserve not only spatial structures within each channel but also the quantitative relationships between channels. This challenge is particularly explicit in frequency-domain FLIM (fdFLIM). In fdFLIM, lifetime is recovered from phase and modulation information encoded in temporally modulated fluorescence signals, typically through ratio-derived relationships between sine and cosine components^20,21^. When fdFLIM is combined with SIM, these modulation channels must be reconstructed from multiple low-SNR raw images. Independent SR reconstruction of the sine and cosine components can introduce subpixel misregistration, alter inter-channel ratios and amplify residual noise during lifetime calculation. As a result, the reconstructed intensity image may appear spatially resolved while the derived lifetime map becomes biased, broadened or unstable. Conventional SIM reconstruction algorithms are designed to restore intensity rather than preserve ratio-derived physical parameters^22,23^, whereas conventional FLIM denoising strategies do not incorporate the optical image-formation model required for SR reconstruction^24,25^. Quantitative SR-FLIM therefore requires a reconstruction framework that jointly considers SR image formation and lifetime estimation.

Here we introduce structured-illumination frequency-domain FLIM (SIM-fdFLIM), a ratio-preserving SR lifetime imaging approach designed to maintain quantitative lifetime fidelity during live-cell SR reconstruction. Rather than treating SIM reconstruction and lifetime estimation as independent steps, SIM-fdFLIM links the optical image-formation model to the ratio-derived lifetime calculation. The method combines direct-current (DC)-stabilized lifetime estimation, which improves the robustness of dual-channel SIM reconstruction and frequency-domain measurements under low-SNR conditions, with **r**atio-domain **i**ntegrated **v**ariational **e**stimation (**RIVE**), which corrects channel misregistration and suppresses ratio-amplified noise after SR reconstruction. We validate this framework using fluorophore standards and simulations designed to separate true lifetime contrast from intensity contrast, and then apply it to live-cell imaging of mitochondrial functional heterogeneity, lipid microenvironment variation and organelle-contact signatures. Together, these results show that preserving ratio-derived quantitative readouts during SR reconstruction enables quantitative functional lifetime imaging at nanoscale resolution in living cells.

## Results

### Principle and the pipeline of SIM-fdFLIM

SIM-fdFLIM integrates spatially structured illumination with widefield frequency-domain FLIM detection using a commercially available high-speed fdFLIM camera (**Fig. 1a**)^21,26,27^. In this system, the SIM excitation pattern is sinusoidally modulated in time, introducing a synchronized spatiotemporal modulation of the fluorescence signal that is jointly sampled by frequency-domain FLIM detection (**Fig. 1b**). The resulting fluorescence signal is recorded by the dual-tap acquisition scheme of the camera (**Fig. 1c**)^21^. Temporal modulation of excitation was implemented using synchronized laser intensity modulation, enabling stable frequency-domain lifetime detection while maintaining high-contrast structured illumination. For each field of view, we acquire nine groups of eight phase-shifted frames (**Supplementary Fig. 1**), each group corresponding to a distinct structured illumination pattern. The recorded signals are decomposed into DC, sine, and cosine components using Fourier analysis, forming the basis for frequency-domain lifetime estimation. Widefield (WF) components are obtained by averaging across the nine illumination patterns, providing a stable reference, whereas pattern-resolved components retain the spatial modulation necessary for SIM reconstruction. To improve robustness in lifetime estimation under low-SNR conditions, we derive a DC-combined lifetime metric that stabilizes widefield lifetime calculation while maintaining consistency with subsequent SIM reconstruction (**Supplementary Note 1, Supplementary Fig. 2)**. Super-resolution intensity images are then reconstructed using Hessian-regularized SIM, enforcing spatiotemporal continuity and suppressing reconstruction artifacts^28^. Finally, we introduce RIVE, which formulates lifetime recovery as a ratio-domain constrained optimization problem. In this framework, a cross-correlation constraint enforces consistency between the cosine- and sine-derived SIM ratio images and the corresponding intensity reconstruction, while a total-variation prior promotes piecewise-smooth lifetime distributions (**Fig. 1d**). Using this complete SIM-fdFLIM pipeline, we first demonstrate that microtubules and mitochondria labeled with spectrally similar fluorophores can be discriminated based on their distinct fluorescence lifetimes, highlighting the modality’s ability to resolve functional contrast beyond intensity imaging (**Fig. 1e-g**). Spatial resolution improvement is quantified using full-width at half-maximum (FWHM) analysis of filament structures, confirming an approximately twofold enhancement from WF to SIM, reaching ∼100 nm lateral resolution (**Fig. 1h**), and positioning SIM-fdFLIM in a favorable speed-resolution regime among representative FLIM modalities (**Fig. 1i**). In lifetime reconstruction, we observe elevated apparent lifetime values in low-signal background regions, which arise from reduced photon counts and the associated decrease in modulation depth in frequency-domain estimation (**Supplementary Note 1, Supplementary Fig. 3)**. To ensure robust quantitative interpretation, subsequent analyses focus on the experimentally relevant lifetime range of 0-4 ns, within which filament width measurements remain consistent with the independently derived ∼100 nm spatial resolution (**Fig. 1h**).

**Fig. 1.**
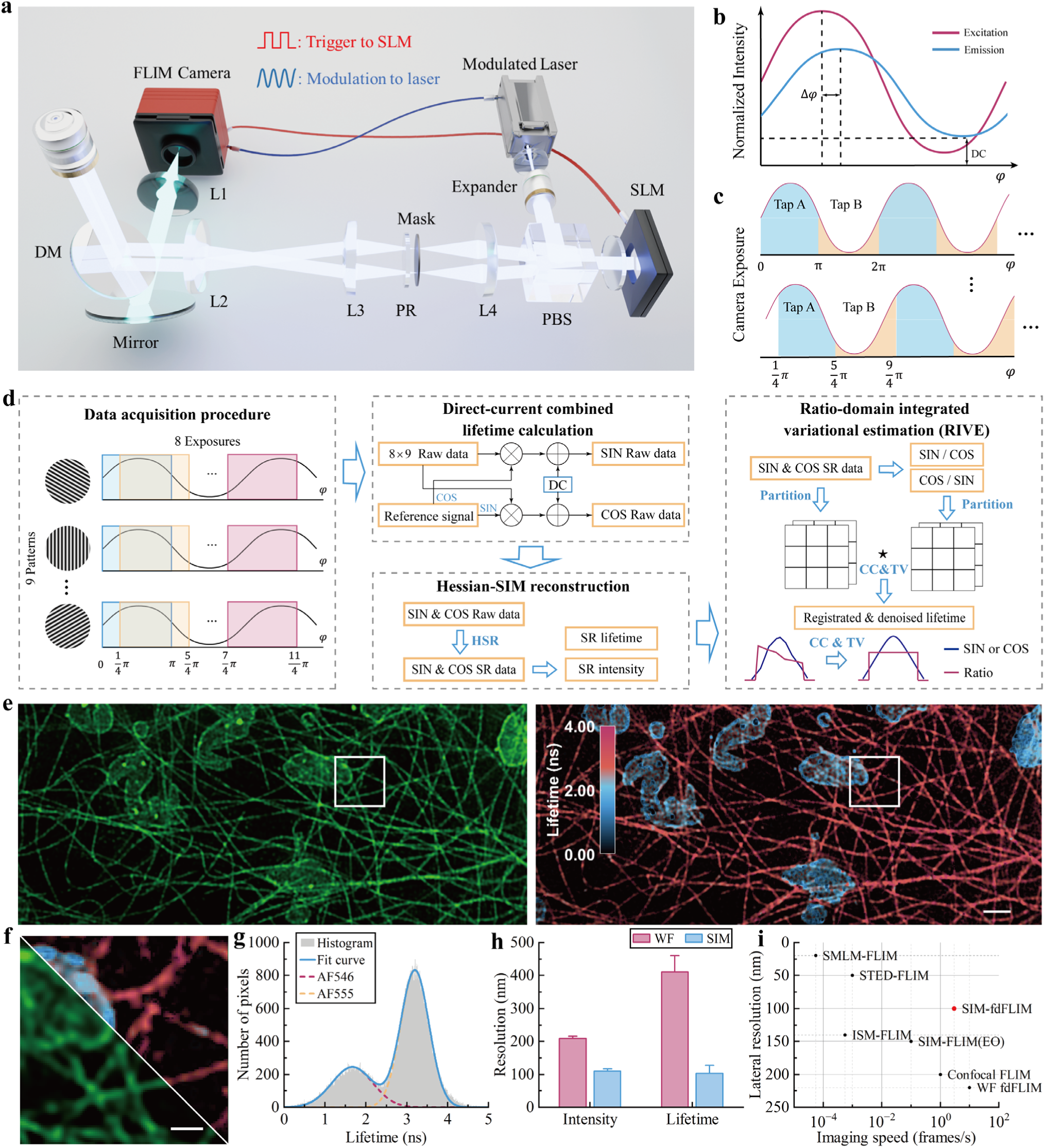
| SIM-fdFLIM system setup and performance. **a,** Schematic of system hardware setup. A detailed illustration is provided in **Methods**. DM, dichroic mirror; L1-L4, lenses; PR, polarization rotator; PBS, polarizing beam splitter; SLM, spatial light modulator. **b,** Temporal modulation of the illumination and the resulting fluorescence emission. The x-axis represents the temporal modulation phase (equivalent to time), and the y-axis represents the normalized illumination intensity at each position in the x-y plane. Δ*φ* denotes the phase delay between excitation and emission associated with fluorescence lifetime. **c**, Camera exposure timing sequence, where the two charge-collection sites (Tap A and Tap B) operate alternately. The x-axis represents the temporal modulation phase (equivalent to time), and the y-axis represents the fluorescence emission. The integrated areas of A and B correspond to the collected signals. **d,** Processing pipeline of SIM-fdFLIM. 9 structured illumination patterns are each acquired over 8 temporal phase shifts, yielding 72 raw images. These data are first processed using DC-combined lifetime estimation to obtain SIN and COS channels for Hessian-SIM reconstruction. RIVE is subsequently applied to the reconstructed SR images to refine lifetime estimation. HSR: Hessian-SIM reconstruction; ∑ : cross-correlation. **e,** Intensity (left) and lifetime (right) images obtained by SIM-fdFLIM. Microtubules were labeled with AF546 and mitochondria were labeled with AF555 in a COS-7 cell. Lifetime image is overlaid with intensity (see **Methods**). **f,** Zoom-in views of the regions indicated in **e**, with intensity images shown at the lower left and lifetime maps at the upper right. **g,** Lifetime histograms corresponding to the lifetime images in **e**, together with double-Gaussian fits. **h,** Bar plots showing resolution in intensity and lifetime for the two imaging systems, presented as mean + SD (*n* = 12). SD: standard deviation. **i,** Trade-off landscape of lifetime-map imaging rate and lateral resolution across representative FLIM modalities. Detailed representative parameters are provided in **Supplementary Table 1**. Scale bar: 1 μm; zoom-in panels: 500 nm.

### DC-combined lifetime calculation

We evaluated the performance of the SIM-fdFLIM system using fixed cells immunofluorescently labeled with antibodies conjugated to a single-lifetime fluorophore, Alexa Fluor 488 (AF488), targeting microtubules. In the fdFLIM detection scheme, sinusoidally modulated fluorescence signals are sampled at multiple phase steps, and relative phase information is extracted to enable frequency-domain lifetime estimation (**Fig. 1c**). For both WF and SIM-fdFLIM, eight phase samples are acquired per modulation cycle, satisfying the Nyquist criterion (≥ 2 points per cycle) for accurate harmonic reconstruction^29^. Frequency-domain analysis of the temporally sampled signals yields the first harmonic components, from which fluorescence lifetime is determined via sine-cosine ratio estimation (**Supplementary Note 1**). However, under temporally modulated excitation, reduced photon flux leads to SNR degradation, which can destabilize the reconstruction of sine and cosine components and introduce artifacts in both widefield and SIM-based ratiometric lifetime maps (**Fig. 2a-e, Supplementary Fig. 2c,f**). To address these problems, we introduce a DC-combined lifetime metric that incorporates the DC component into both sine- and cosine-channel representations, yielding a monotonic and more noise-robust formulation of the ratio-based lifetime estimator (**Supplementary Note 1, Supplementary Fig. 2a,b)**. The DC-combined sine and cosine channels are denoted as SIN and COS, respectively. Importantly, because both DC and first-harmonic components remain linearly related to the underlying modulated fluorescence signal, this formulation preserves the modulation structure required for subsequent SIM reconstruction without altering spatial modulation depth (**Fig. 1d**, **Supplementary Fig. 2a** and **Supplementary Note 1**). This DC-combined formulation significantly improves reconstruction robustness, increasing the fraction of successfully reconstructed SIM images and enhancing SIN-channel SNR by ∼2 dB (∼1.58-fold, **Fig. 2c, Supplementary Fig. 2d**). To directly quantify performance in the photon-limited regime, we binned repeated measurements according to the background-subtracted sine-component intensity, used as a proxy for the detected photon budget. As shown in **Fig. 2e**, when referenced to the WF-fdFLIM lifetime distribution shown in **Fig. 2d**, conventional sine-cosine lifetime estimation exhibited increasing lifetime deviation and frame-to-frame drift as signal decreased, whereas the DC-combined formulation maintained mean lifetimes closer to the WF reference and substantially reduced lifetime variability across the same signal range. These results demonstrate that DC-combined preprocessing improves the photon-budget tolerance of SIM-fdFLIM during SR lifetime reconstruction under the same RIVE processing conditions (**Supplementary Fig. 2g-h**).

**Fig. 2.**
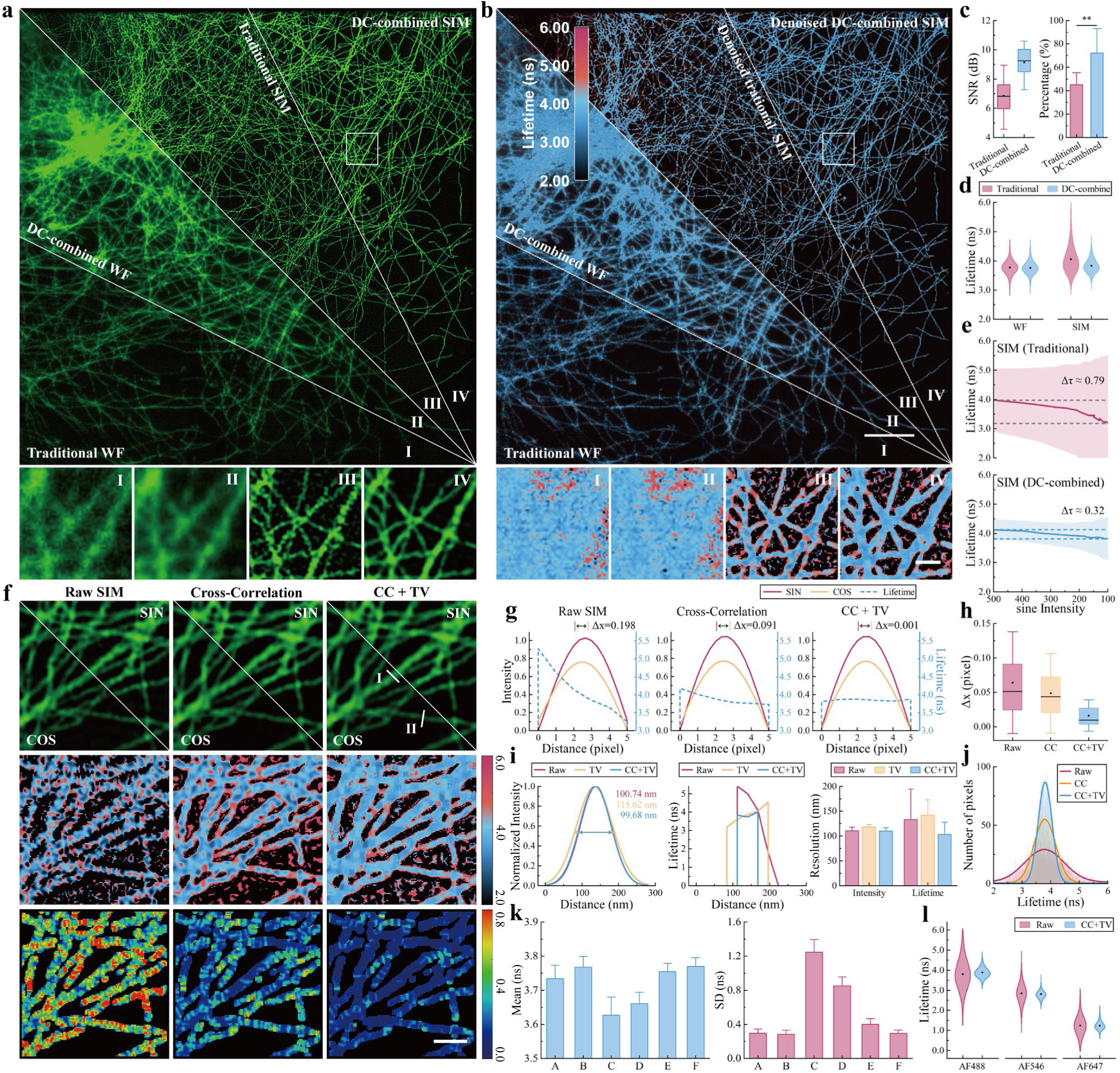
| SIM-fdFLIM stabilizes ratio-derived lifetime estimation and channel registration. **a,** Intensity images of the SIN channel obtained by traditional WF-fdFLIM (I), DC-combined WF-fdFLIM (II), Wiener SIM reconstruction from I (III) and II (IV), respectively. Scale bar: 10 μm; zoom-in panels: 500 nm. **b,** Lifetime images overlaid with SIN-channel intensity images, obtained by traditional WF-fdFLIM (I), DC-combined WF-fdFLIM (II), and SIM-fdFLIM reconstructed by Hessian-SIM with RIVE (denoised SIM-fdFLIM) from I (III) and II (IV), respectively. The zoom-in panels below show the pure lifetime images without intensity overlay. Scale bar: 10 μm; zoom-in panels: 500 nm. **c**, Box plots showing the signal-to-noise ratio (SNR) of traditional and DC-combined WF intensity images (left). Bar plots showing percentage of traditional and DC-combined SIM reconstructions with variance below 0.5 (see **Methods**), presented as mean + SD (right, *n* = 12 captures per group). Data are from SIN channel and presented as mean (dot), median (center line), interquartile range (box), and 5-95% range (whiskers). *n* = 12 regions of interest (ROIs) per group. **d,** Violin plots showing lifetime histograms of traditional ( μ = 3.77; σ = 0.21) and DC-combined ( μ = 3.75; σ = 0.19) WF-fdFLIM (left), and traditional ( μ = 3.95; σ = 0.41) and DC-combined ( μ = 3.79; σ = 0.23) denoised SIM-fdFLIM (right). μ: mean; σ: SD. Data were derived from the images shown in **b**. **e,** Photon-budget-dependent lifetime performance of SIM-fdFLIM. Lifetime measurements obtained using traditional (top) and DC-combined (bottom) denoised SIM-fdFLIM are plotted as a function of background-subtracted average SIN-channel intensity, used as a proxy for detected photon budget. The WF-fdFLIM lifetime distribution in **d** serves as the reference for evaluating signal-dependent lifetime deviation. Δτ denotes frame-to-frame drift in mean lifetime. Shaded regions represent SD, *n* ≥ 12 ROIs. **f,** Zoom-in views of **a** showing SIM reconstructions without RIVE (Raw SIM-fdFLIM, termed Raw SIM; left column), SIM with CC regularization (middle column), and SIM with CC + TV regularization (right column). All SIM results are reconstructed from DC-combined widefield data. Top row: intensity images; middle row: pure lifetime images; bottom row: local standard deviation maps of lifetime. Scale bar: 500 nm. **g,** Line profiles along the lines I in **f**. The intensity profile was normalized and fitted with a Gaussian function. Δ *x* indicates the distance between the centroid positions of Gaussian fits in the SIN and COS channels. Improved flatness of lifetime distribution is observed as this distance decreases. **h,** Box plots showing the centroid distance under different regularization strategies. Data presentation is the same as in **c**. *n* = 24 ROIs per group. **i,** Line profiles along the lines II in **f** indicating the spatial resolution in intensity (normalized Gaussian-fitted, left) and lifetime (middle) for different regularization strategies. Bar plots on the right show the statistical results, presented as mean + SD (*n* = 12). **j,** Lifetime histograms and the corresponding Gaussian fits of SIM-fdFLIM images reconstructed using different regularization strategies. **k,** Mean (left) and SD (right) of lifetime values measured from AF488-labeled microtubule samples across repeated acquisitions. Bar plots are presented as mean + SD (*n* = 12). A: traditional WF-fdFLIM; B: DC-combined WF-fdFLIM; C: traditional SIM-fdFLIM without RIVE; D: DC-combined raw SIM-fdFLIM without RIVE; E: DC-combined SIM-fdFLIM w/ CC regularization; F: DC-combined SIM-fdFLIM w/ CC+TV regularization. **l,** Violin plots showing fluorescence lifetime distributions of raw SIM-fdFLIM and DC-combined SIM-fdFLIM with CC+TV regularization for different immunofluorescence samples (AF488, AF546, and AF647).

### Ratio-domain instability in SIM-fdFLIM and RIVE-based reconstruction

Following DC-combined preprocessing, SIM images are reconstructed using the Hessian-SIM reconstruction algorithm, which incorporates spatial and temporal continuity priors to reduce random artifacts induced by noise amplification^28^. However, directly computing lifetimes from the reconstructed SIN/COS pair introduces two failure modes. First, independent reconstruction of SIN and COS images can introduce subpixel misregistration, which minimally affects intensity but produces pronounced artifacts after ratiometric division and lifetime conversion (**Fig. 2f-h**, **Supplementary Fig. 5**). This effect originates from statistical uncertainty in PSF center estimation, which can be formally described by the Cramér-Rao lower bound (CRLB)^30,31^. Although this uncertainty is typically negligible in intensity-based imaging at ∼100 nm resolution, it becomes critical in ratio-based estimation due to the sensitivity of ratiometric operations to spatial misalignment. Second, ratio formation amplifies residual noise, broadening lifetime distributions and obscuring separation of multiple lifetime populations (**Fig. 2j-l, Supplementary Fig. 6-9**). This behavior is consistent with classical error propagation theory^32^, which predicts that variance is significantly amplified under nonlinear division operations. Taken together, these two mechanisms reveal an intrinsic inconsistency between super-resolution reconstruction and ratio-based lifetime estimation, motivating a reconstruction framework that operates directly in the ratio domain to ensure estimator consistency and stable lifetime recovery. In the following analyses, SIM reconstructions without the RIVE step are referred to as Raw SIM-fdFLIM (Raw SIM), and Hessian-SIM reconstructions with RIVE are referred to as denoised SIM-fdFLIM (denoised SIM). Where indicated, Wiener-SIM reconstruction of the SIN and COS channels followed by direct ratio conversion was used as the non-RIVE baseline.

We formulate RIVE as a multiplicative iterative optimization framework with joint regularization (**Eq. (13)** in **Supplementary Note 2**). Two symmetric cross-correlation (CC) regularization terms enforce structural consistency between the intensity and ratio images, correcting subpixel mismatches that would otherwise reduce their correlation (**Fig. 2f-h**) and suppressing background artifacts in the reconstructed lifetime maps. In the absence of CC regularization, neither total variation (TV) nor Hessian priors can fully correct spatial misalignment, leading to reduced intensity-lifetime agreement and increased lifetime bias (**Supplementary Fig. 5**).

To further stabilize lifetime estimation under noise, a TV prior is incorporated to promote piecewise-smooth lifetime distributions while preserving structural boundaries. CC alone cannot suppress spatially heterogeneous lifetime fluctuations, even in samples labeled with single-lifetime fluorophores (**Supplementary Fig. 6**), whereas TV alone leads to oversmoothing of structural details and reduced spatial resolution (**Fig. 2i, Supplementary Fig. 6d,e**), biased lifetime distributions (**Supplementary Fig. 6g**), and diminished contrast in discriminating multiple lifetime species (**Supplementary Fig. 8**). The combination of CC and TV resolves these limitations by enforcing spatial consistency while constraining lifetime variability. In practice, although CC+TV only marginally affects spatial resolution as assessed by FWHM and decorrelation analysis (DCA) of intensity images (**Fig. 2i, Supplementary Fig. 6d-f**), it substantially improves the fidelity of lifetime distributions (**Supplementary Fig. 6g**) and the separation of multiple lifetime populations (**Supplementary Fig. 8**).

Therefore, minimizing the combined CC and TV penalty effectively compensates for random noise and spatial mismatches, and suppresses background contributions. With only minimal perturbations introduced to intensity values, SR structures and realistic lifetime variations are preserved (**Supplementary Fig. 5-9**). Compared with Raw SIM-fdFLIM^22,33^, RIVE-reconstructed SIM-fdFLIM yielded substantially narrower Gaussian lifetime distributions for microtubules, with mean lifetimes closely matching WF-fdFLIM. Across AF488-, AF546-, and AF647-labeled single-lifetime samples spanning 1.24–3.78 ns, RIVE reduced lifetime standard deviations by 2.5- to 3-fold relative to Raw SIM-fdFLIM (**Fig. 2k-l, Supplementary Fig. 7**). Simulations further showed that objects with three distinct ratios, each directly corresponding to a different lifetime, became indistinguishable in the presence of reconstruction noise but were clearly separated after RIVE processing. This ratio contrast remained stable across intensity conditions, including structures with identical intensities but different ratios and structures with different intensities but identical ratios (**Supplementary Fig. 9**). These controls indicate that RIVE preserves ratio-derived contrast rather than transferring structural intensity contrast into the lifetime map. Overall, RIVE enables SIM-fdFLIM to generate high-resolution, low-noise and quantitatively faithful lifetime maps by simultaneously reducing reconstruction-induced bias and noise-induced variance, while preserving the integrity of super-resolution spatial information (**Supplementary Note 2**).

### Rapid SIM-fdFLIM reveals hierarchical mitochondrial heterogeneity

Mitochondrial membrane potential (ΔΨm) can be reported by lifetime-sensitive probes, which are less sensitive to illumination nonuniformity and other intensity fluctuations than intensity-only dyes. PK Mito Deep Red (PKMDR) was recently introduced as a mitochondrial lifetime probe with concentration-dependent lifetime behavior, enabling dual validation through intensity and lifetime readouts^34^. Within the same probe family, we found that PK Mito Red (PKMR) exhibits similar intensity-lifetime characteristics to PKMDR, consistent with their structural similarity. Owing to its shorter emission wavelength, PKMR provides higher spatial resolution and is well suited for resolving fine mitochondrial structures such as cristae.

Combined with SIM-fdFLIM, PKMR enabled clear visualization of cristae together with SR lifetime mapping (**Fig. 3a**). SIM-fdFLIM revealed a pronounced negative correlation between fluorescence intensity and lifetime: intensity maxima aligned with cristae and coincided with local lifetime minima, whereas this relationship was largely obscured in WF-fdFLIM (**Fig. 3b,c**). Increased PKMR accumulation elevated fluorescence intensity while shortening lifetime, consistent with spatial variation in a probe-reported, membrane-potential-linked mitochondrial state at the scale of cristae. The stability of lifetime profiles across frames further excludes reconstruction artifacts. Notably, the cross-correlation between adjacent frames is higher for SIM-fdFLIM than for WF-fdFLIM, with the lower values in WF-fdFLIM attributable to noise (**Fig. 3d**). This behavior parallels prior observations obtained by intensity-based membrane potential dyes such as TMRE, but SIM-fdFLIM enables this contrast to be quantified directly in lifetime space at sub-organelle resolution^35^.

**Fig. 3.**
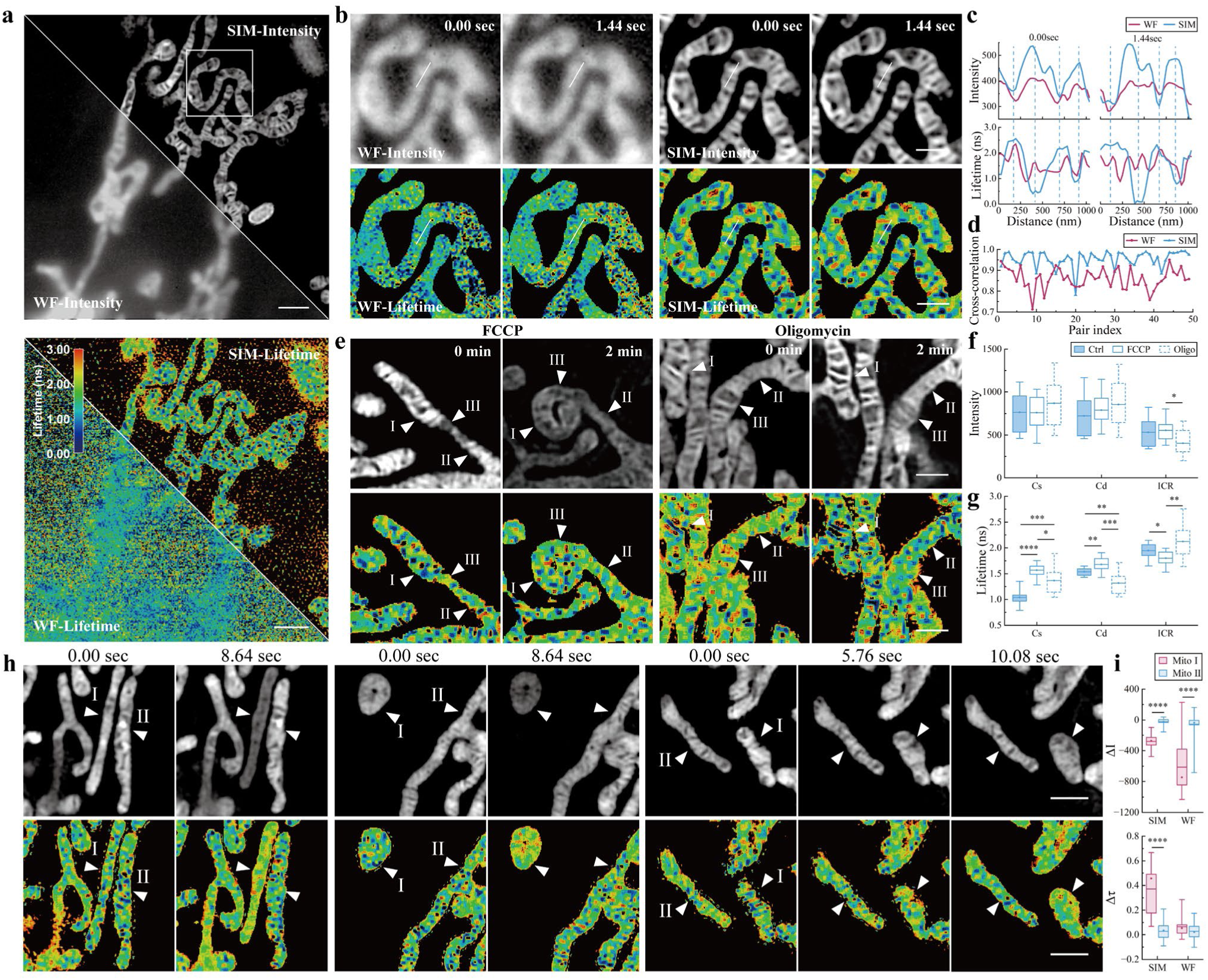
| Live cell lifetime imaging of mitochondria stained with PK Mito Red (PKMR). **a,** Comparison of intensity images obtained by SIM and widefield (WF) imaging (top), and the corresponding lifetime distributions measured by SIM-fdFLIM and WF-fdFLIM (bottom). Scale bar: 3 μm. **b,** Zoom-in time-lapse images from **a** showing the distribution of mitochondrial inner membrane intensity (top row) and lifetime (bottom row) acquired by WF (left) and SIM (right). Scale bar: 1 μm. **c,** Quantified intensity and lifetime profiles along the lines indicated in **b**, revealing an inverse relationship between intensity and lifetime. **d,** Cross-correlation curves between lifetime profiles of adjacent frame pairs in **b**, supporting temporal consistency of reconstructed lifetime profiles. Each frame has a duration of 1.44 s. **e,** Intensity (top row) and lifetime (bottom row) images of mitochondria treated with FCCP or oligomycin from SIM-fdFLIM. Images at 0 min represent untreated controls. Arrow I: sparse cristae; II: dense cristae; III: inter-cristae regions. Scale bar: 1 μm. **f,** Box plots showing the mean intensity of mitochondrial internal structures under different conditions. Cs: sparse cristae; Cd: dense cristae; ICR: inter-cristae region. *n* ≥ 54 Cs, *n* ≥ 54 Cd, and *n* ≥ 36 ICRs from ≥ 10 cells. **g,** Box plots showing the mean lifetime of mitochondrial internal structures under different conditions. *n* ≥ 12 cells. **h,** Representative mitochondria exhibiting rapid changes in intensity (top row) and lifetime (bottom row) from SIM-fdFLIM. I: changing mitochondria; II: stable mitochondria. Left: tubular changing mitochondria; middle: round changing mitochondria; right: tubular-to-round transitioning changing mitochondria. Scale bar: 1 μm. **i,** Box plots showing the variation of intensity (top) and lifetime (bottom) during mitochondrial changes. For visualization purposes only, background pixels outside the intensity mask and pixels with lifetimes greater than 3 ns were set to black; all quantitative analyses were performed using the indicated intensity masks (except in **a**). The unmasked version is provided in **Supplementary** Fig. 10. For all box plots, data are presented as mean (filled circle), median (center line), interquartile range (box), and 5-95% range (whiskers). For all bar plots, data are presented as mean with positive SD error bars. *P < 0.05; **P < 0.01; ***P < 0.001; ****P < 0.0001.

Closer inspection further revealed heterogeneity across cristae microdomains. We distinguished cristae-dense (Cd) regions, containing multiple closely spaced cristae that are not resolvable at ∼100 nm resolution, from cristae-sparse (Cs) regions that are spatially separated from neighboring folds (**Fig. 3e-g**). Cs regions exhibited the shortest lifetimes and highest intensities, whereas Cd regions showed intermediate lifetimes relative to isolated cristae and the surrounding mitochondrial interior, with intensities comparable to those of isolated cristae. In contrast, inter-cristae regions (ICRs) exhibited the longest lifetimes and lowest intensities (**Fig. 3f,g**). Pharmacological perturbations supported a functional origin of these differences. FCCP, which dissipates the proton gradient, increased lifetimes at both cristae-dense and cristae-sparse sites while decreasing lifetime in the ICRs. In contrast, oligomycin produced a differential response: lifetimes increased at Cs segments but decreased at Cd regions, with minimal change in the surrounding interior (**Fig. 3g**). Because FCCP and oligomycin perturb mitochondrial proton-motive force through distinct mechanisms, their divergent effects further support the functional nature of cristae-scale lifetime heterogeneity. Given that PKMR-family probes report mitochondrial membrane-potential-linked dye accumulation^34^, we interpret these shifts as local functional signatures of mitochondrial state rather than an absolute calibration of ΔΨm. Together, these data indicate that cristae-associated microdomains respond differently to perturbations of proton motive force and demonstrate that SIM-fdFLIM resolves functional variation across cristae-scale mitochondrial substructures.

The high imaging rate of SIM-fdFLIM further enables continuous tracking of mitochondrial dynamics in live cells. At enhanced spatial resolution, we identified a subset of individual mitochondria (type I)—including tubular (∼50%), round (∼40%), and tubular-to-round transitioning (∼10%)—exhibiting rapid intensity loss accompanied by a concomitant increase in lifetime over ∼8–10 s, while neighboring mitochondria (type II) within the same cell remained comparatively stable (**Fig. 3h**). In total, 13 type I mitochondrial events were identified across 13 cells. Interestingly, lifetime differences between type I and type II mitochondria were most apparent in the SIM-fdFLIM measurements and were largely absent in WF-fdFLIM, despite clear intensity differences in both WF and SIM images (**Fig. 3i, Supplementary Fig. 10**). This contrast highlights the ability of SIM-fdFLIM to uncover subtle functional heterogeneity that is obscured at diffraction-limited resolution. Similar mitochondrial heterogeneity has been reported using PKMDR, but limited spatial and temporal resolution confined those observations to whole-cell averages at ∼0.1 Hz^34^. Overall, these results indicate that mitochondria within a single cell—and cristae within individual mitochondria—can form distinct functional subpopulations with divergent intensity and lifetime trajectories, highlighting the ability of SIM-fdFLIM to measure dynamic energetic heterogeneity across hierarchical mitochondrial structures. While the profile-stability analysis in **Fig. 3b-d** used 1.44-s frame intervals, faster mitochondrial dynamics, including fission, fusion and nanotunnel formation, were recorded at 2.8 Hz under optimized PKMR imaging conditions (**Supplementary Video 4**).

### SIM-fdFLIM resolves sub-organelle lipid heterogeneity and organelle contact microenvironments

Nile Red is a well-established reporter of membrane polarity, viscosity, and lipid order^36,37^. Previous studies have used Nile Red to label membrane structures across distinct subcellular compartments, with segmentation primarily based on organelle morphology^38,39^. Although Nile Red lifetime and spectral readouts have been used to report intracellular lipid environments^40,41^, prior studies have not combined live-cell-compatible widefield acquisition, super-resolution lifetime reconstruction, and simultaneous analysis of organelle-specific and contact-site membrane microenvironments.

In live COS-7 cells stained with Nile Red, SIM-fdFLIM resolved nanoscale membrane architectures, including the endoplasmic reticulum (ER), mitochondria (MITO), lipid droplets (LD), and the plasma membrane-dominant peripheral regions (MEM), each exhibiting distinct lifetime characteristics (**Fig. 4a-c**). However, the global lifetime distribution pooled from all fluorescence-emitting pixels can be biased by the relative abundance and size of different subcellular structures within the field of view. To address this, we segmented pixels based on organelle morphology— tubular networks for ER, cristae-rich regions for MITO, spherical droplets for LD, and peripheral membrane regions for MEM—prior to normalizing and jointly plotting their lifetime distributions. This analysis yielded four partially overlapping yet distinct Gaussian lifetime distributions with graded mean values: LD exhibited the shortest lifetime (2.74 ± 0.31 ns), followed by MITO (2.98 ± 0.29 ns), ER (3.42 ± 0.25 ns), and MEM (4.00 ± 0.51 ns) (**Fig. 4b**). In contrast, under WF lifetime imaging, statistically significant lifetime differences were only observed for LD (3.02 ± 0.19 ns) and MEM (3.22 ± 0.29 ns), whereas MITO and ER were not distinguishable (**Supplementary Fig. 11b**). Consistent with this, WF measurements showed substantially greater overlap among organelle-specific lifetime distributions, and the mean lifetime separation between LD and MEM was reduced to approximately one-third of that achieved with SIM-fdFLIM. Pharmacological perturbations further validated the environmental origin of the observed lifetime contrast. Paclitaxel (PTX) treatment reduced the fluorescence lifetimes of all four organelles to varying extents under SIM-fdFLIM, whereas these lifetime reductions were largely absent in WF measurements. In contrast, oleic acid (OA) increased the lifetime of LD by ∼0.2 ns under both SIM and WF modes, while selectively elevating the MEM lifetime in WF imaging. Following OA treatment, the lifetime differences among LD, MITO, and ER became less pronounced, consistent with a homogenization of membrane environments (**Fig. 4b,c, Supplementary Fig. 11b,c**).

**Fig. 4.**
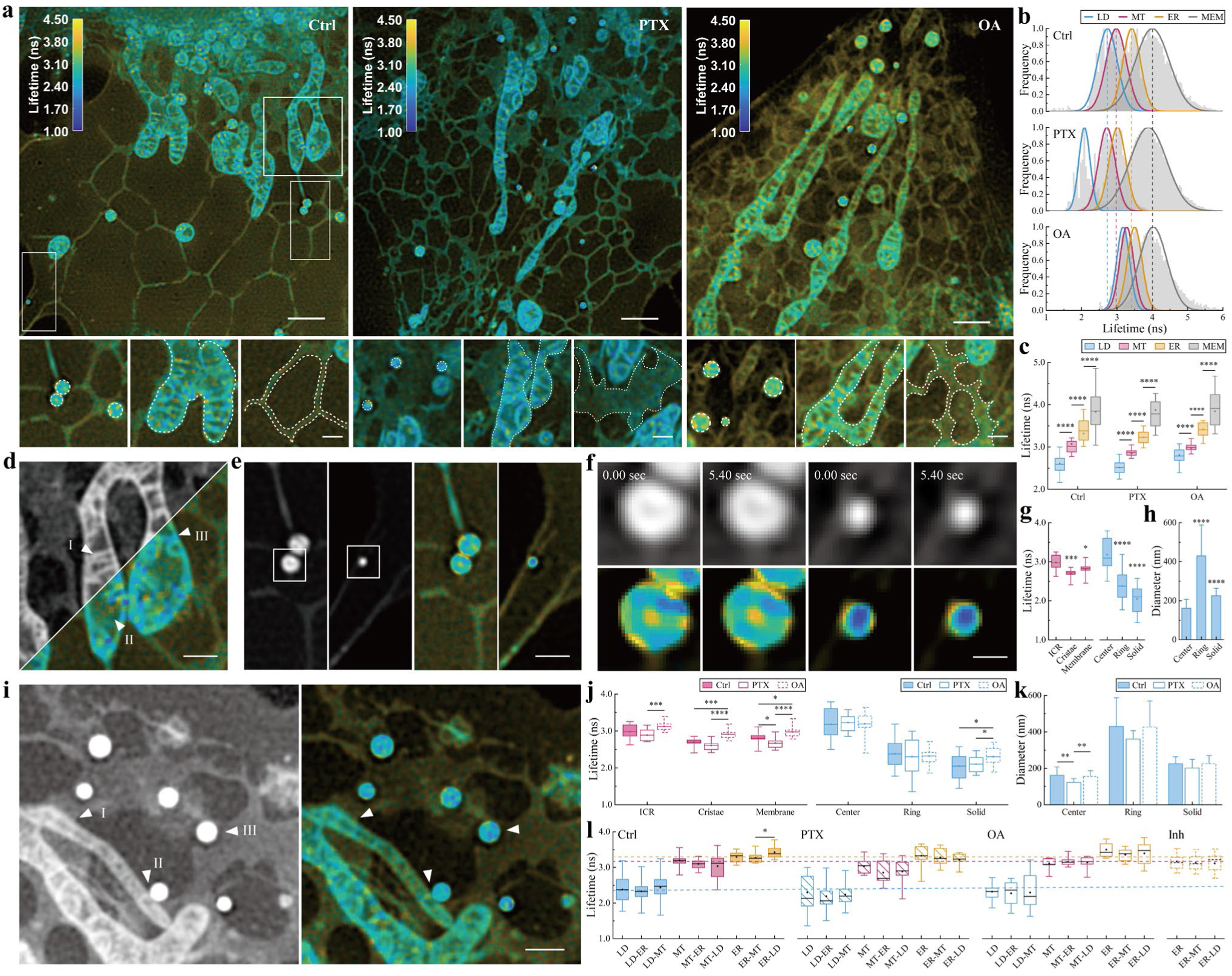
| SIM-fdFLIM imaging of live cells stained with Nile Red. **a,** Representative intensity-weighted lifetime images of untreated (left, Ctrl), paclitaxel-treated (middle, PTX), and oleic acid–treated (right, OA) cells. The bottom row shows magnified views of lipid droplets, mitochondria and endoplasmic reticulum for each condition, with dashed outlines indicating ROIs manually segmented based on organelle morphology. Scale bar: 2 μm; zoom-in panels: 500 nm. **b,** Lifetime distributions and the corresponding Gaussian fits of lipid droplets (LD), mitochondria (MITO), endoplasmic reticulum (ER) and the plasma membrane-dominant peripheral regions (MEM) under the conditions shown in **a**. **c,** Box plots showing the distributions of mean lifetimes of LD, MITO, ER, and MEM under different conditions. *n* ≥ 34 ROIs. **d,** Intensity (upper-left) and intensity-weighted lifetime (lower-right) images of mitochondria from the selected regions of interest in a. Triangle arrows indicate the cristae (I), ICR (II), and membrane (III) regions of mitochondria. Scale bar: 500 nm. **e,** Intensity (left) and intensity-weighted lifetime (right) images of ring-shaped and solid lipid droplets, arranged horizontally. Scale bar: 500 nm. **f,** Intensity (top row) and intensity-weighted lifetime (bottom row) images of the lipid droplets indicated in e, shown at consecutive frames (5.4 s apart). Scale bar: 200 nm. **g,** Box plots showing the distributions of mean lifetimes of internal structures in mitochondria (left) and lipid droplets (right). *n* ≥ 12 cells. **h,** Bar plots showing the diameters of solid lipid droplets and the inner and outer diameters of ring-shaped lipid droplets. *n* ≥ 12 cells. **i,** Intensity (left) and Intensity-weighted lifetime (right) images of organelle contact regions. Triangle arrows indicate the contact regions between mitochondria and endoplasmic reticulum (I), mitochondria and lipid droplets (II), and endoplasmic reticulum and lipid droplets (III). Scale bar: 500 nm. **j,** Box plots showing the distributions of mean lifetimes of internal regions in mitochondria (left) and lipid droplets (right) under different conditions. *n* ≥ 12 cells. **k,** Bar plots showing the diameters of internal regions in lipid droplets under different conditions. *n* ≥ 12 cells. **l,** Box plots showing mean lifetimes quantified within 112-nm-diameter ROIs centered on the indicated contact-side regions. Inh, combined DGAT1 and DGAT2 inhibition. Labels: A denotes the bulk region of organelle A, whereas A-B denotes the contact region on the A side of the A-B interface. *n* ≥ 12 cells. For all box plots, data are presented as mean (filled circle), median (center line), interquartile range (box), and 5-95% range (whiskers). For all bar plots, data are presented as mean with positive SD error bars. *P < 0.05; **P < 0.01; ***P < 0.001; ****P < 0.0001.

Leveraging the SR capability of SIM-fdFLIM, we further resolved nanoscale lifetime heterogeneity within individual organelles, including individual mitochondria and LDs (**Fig. 4d-f**). Within mitochondria, the ICRs exhibited significantly longer lifetimes than both the cristae and the membrane, whereas the latter two were indistinguishable from each other (**Fig. 4g**); this ICR-associated lifetime prolongation was not observed with WF-fdFLIM (**Supplementary Fig. 11i**). PTX treatment modestly reduced lifetimes across all mitochondrial subcompartments, reaching statistical significance at the membrane. In contrast, OA treatment generally increased lifetimes relative to PTX and elevated the membrane lifetime above control levels (**Fig. 4j**). Notably, we identified two distinct populations of LDs: larger droplets with a mean diameter of ∼430 nm that exhibited a Nile Red-depleted core, and smaller droplets with a homogeneous core and a mean diameter of ∼225 nm (**Fig. 4e-h**). The larger LDs displayed pronounced intradroplet lifetime heterogeneity, with significantly longer lifetimes at the core than at the periphery or those of the smaller, homogeneous LDs (**Fig. 4f,g**); these differences were likewise absent in WF-fdFLIM (**Supplementary Fig. 11i**). Whereas PTX treatment did not significantly affect the lifetimes of either large or small LDs, OA treatment selectively increased the lifetime of small LDs (**Fig. 4j**), consistent with the population-level analysis of all lipid droplets (**Fig. 4c**). Together, these results reveal previously inaccessible nanoscale variations in membrane and lipid environments within individual organelles.

Finally, we examined fluorescence lifetimes at contact sites between the ER, MITO, and LDs (**Fig. 4i,l**). Despite active lipid exchange between these organelles^40,42^, most contact sites did not exhibit lifetimes distinct from those of the corresponding bulk organelles or other contact regions. Under control conditions, ER-LD contacts displayed a distinct high-lifetime state relative to ER-MITO contacts and non-contact ER regions (**Fig. 4l**), which was also not observed with WF-fdFLIM (**Supplementary Fig. 11k**). Distance-profile analysis further revealed spatially restricted lifetime changes around organelle contacts: relative to bulk ER, ER regions proximal to LDs showed a peak lifetime increase at ∼224 nm from the LD boundary, whereas more distal ER regions showed reduced lifetimes. In contrast, ER regions located 56-224 nm from mitochondria showed elevated lifetime relative to bulk ER (**Supplementary Fig. 12**). This contact-specific lifetime elevation was attenuated by PTX treatment, OA loading and DGAT inhibition, suggesting that the ER-LD high-τ state depends on lipid metabolic flux and preserved subcellular organization (**Fig. 4l**). Thus, SIM-fdFLIM reports a functional state of ER-LD contacts rather than contact proximity alone. Collectively, these results demonstrate that SIM-fdFLIM enables the detection of subtle, spatially restricted heterogeneity in membrane lipid environments at organelle contact sites, revealing context-dependent biophysical properties that are not readily resolved by conventional WF-fdFLIM.

## Discussion

In this work, we establish SIM-fdFLIM as a ratio-preserving strategy for quantitative SR fluorescence lifetime imaging in living cells. The key advance is not only the combination of SIM and FLIM, but the preservation of ratio-derived lifetime information during photon-limited SR reconstruction. By combining DC-stabilized lifetime estimation with RIVE, SIM-fdFLIM achieves ∼100 nm lifetime maps at Hz-rate live-cell imaging while maintaining mean lifetime fidelity and reducing reconstruction-induced lifetime variance. This combination of nanoscale spatial resolution, live-cell temporal resolution and quantitative lifetime preservation defines the main performance advance of the method.

Compared with point-scanning SR-FLIM methods such as STED-FLIM and ISM-FLIM, SIM-fdFLIM benefits from parallel acquisition and therefore offers a favorable balance among temporal resolution, field of view and illumination dose (**Supplementary Table 1**). Recent EO-SIM-FLIM demonstrated live-cell SR lifetime imaging, but SIM-fdFLIM differs by using commercial dual-tap fdFLIM detection and by explicitly correcting ratio-domain artifacts arising from channel misregistration and noise amplification. This design lowers the barrier to implementation while placing SIM-fdFLIM in a performance regime optimized for dense, dynamic live-cell functional imaging.

This ratio-preserving principle is not limited to fdFLIM. Many functional optical measurements generate parameter maps by merging or ratioing multiple images acquired from different spectral channels, temporal gates, polarization states or excitation conditions, including fluorescence resonance energy transfer (FRET)^43^, ratiometric ion indicators^44^, polarization anisotropy^45^ and spectral unmixing^46^. In these modalities, independent denoising or SR reconstruction of each channel can distort the derived physical parameter even when individual images appear visually improved. By formulating reconstruction in the parameter-generating ratio domain, SIM-fdFLIM suggests a potentially generalizable computational strategy for extending functional microscopy beyond the diffraction limit while maintaining quantitative interpretability.

Using the lifetime-sensitive mitochondrial probe PK Mito Red, SIM-fdFLIM resolved membrane-potential-linked lifetime heterogeneity at multiple hierarchical levels, from cristae-associated microdomains to mitochondria-to-mitochondria subpopulations within the same cell. The ability to simultaneously resolve cristae-level microdomains and whole-mitochondrion subpopulations highlights a hierarchical organization of mitochondrial function that is not accessible with diffraction-limited FLIM or purely structural SR methods, while also enabling dynamic tracking in living cells. We note that the system’s effective focal depth (∼400 nm) is smaller than typical mitochondrial thickness (∼700 nm), so out-of-focus contributions may influence lifetime values in intermediate regions (**Supplementary Note 3**); intensity images should therefore serve as a reference for identifying reliably in-focus structures. These considerations do not alter the main conclusions, but underscore the importance of using intensity images and structural context when interpreting lifetime values in axially overlapping regions.

Using Nile Red, SIM-fdFLIM further resolved nanoscale lifetime differences among organelles and within individual mitochondria and lipid droplets. Nile Red lifetime reflects a composite of polarity, hydration, microviscosity, protein crowding and dye partitioning, rather than a single membrane-order parameter. Therefore, the observed lifetime maps should be interpreted as microenvironmental signatures. Sub-organelle lipid heterogeneity has previously been demonstrated using SR intensity-, spectral-, or polarity-based approaches^39,47,48^; however, SIM-fdFLIM extends these observations by providing a lifetime-based, quantitative readout that is less sensitive to illumination nonuniformity and enables direct comparison across structures at SR. Within this framework, the ability to separate lipid droplets, mitochondria, endoplasmic reticulum and plasma membrane, and to resolve lifetime heterogeneity within lipid droplets and mitochondria, demonstrates the utility of SIM-fdFLIM for mapping nanoscale chemical environments in living cells. A particularly informative application is the analysis of organelle contact sites. At ER-LD contacts, SIM-fdFLIM revealed contact-associated microenvironmental states that were not resolved by WF-fdFLIM. More broadly, this example shows how SR lifetime imaging can move beyond structural contact detection toward measurement of contact-associated chemical and biophysical states.

Several limitations remain. SIM-fdFLIM requires multiple temporally and spatially modulated raw frames for each reconstructed lifetime map, which limits temporal resolution. The current implementation is also based on 2D-SIM and widefield detection, making it sensitive to out-of-focus fluorescence and axial overlap in thick structures. In addition, the DC-stabilized lifetime metric assumes single-component decay; in mixed-lifetime regions, the recovered value represents a stable weighted intermediate rather than a full multicomponent decomposition. Future developments combining optical sectioning, 3D-SIM, adaptive illumination, improved camera sensitivity or model-based multicomponent fitting could further extend SIM-fdFLIM toward volumetric and faster functional SR imaging.

Together, these results establish SIM-fdFLIM as a quantitative SR-FLIM approach that preserves ratio-derived lifetime information during live-cell SR reconstruction. By combining nanoscale spatial resolution, Hz-rate imaging and preserved lifetime fidelity, SIM-fdFLIM provides a general route for mapping molecular and microenvironmental states in living cells beyond the diffraction limit.

## Methods

### Optical setup

The SIM-fdFLIM system integrated SIM with widefield fdFLIM (**Fig. 1a**). Fluorescence lifetime detection was performed using a commercially available dual-tap fdFLIM camera (PCO.FLIM X, PCO, 1008 × 1008 pixels, 5.6 μm), where each pixel contained two charge-collection sites (A and B taps) that alternately sample the temporally modulated fluorescence signal. By introducing defined phase offsets between successive capture pairs, the effective temporal sampling rate can be increased (**Fig. 1c**).

Although the Nyquist criterion requires at least two samples per modulation period to avoid aliasing, robust estimation of the DC, sine and cosine components requires additional phase samples. We therefore acquired eight phase-shifted frames for each illumination pattern to improve lifetime precision with dual-tap detection. The excitation laser is modulated both temporally and spatially. Two-dimensional SIM reconstruction requires illumination patterns with three orientations and three phase shifts; accordingly, each SIM-fdFLIM acquisition consisted of nine structured illumination patterns, each paired with eight phase-resolved lifetime frames, yielding a total of 72 raw images per reconstructed frame.

Multicolor excitation was implemented using modulated lasers at 488 nm, 561 nm and 638 nm (Multi-LASER, Lambert Instruments), with modulation frequencies selectable from 0 to 40 MHz. A frequency of 20 MHz was applied consistently across all samples. The output lasers were then collimated and expanded by an objective lens (Expander, CFI Plan Apo 2× objective, NA = 0.10, Nikon). High-contrast structured illumination was generated using a spatial light modulator (SLM; SXGA-3DM or QXGA-3DM, Fourth Dimension Display), a half-wave plate (HWP), and a homemade partitioned polarization rotator (PR)^28^. The modulated beam was routed through a polarizing beam splitter and a 4f relay (ACT508-200-A, Thorlabs), with a spatial filter placed at the Fourier plane to isolate the ±1 diffraction orders. A second 4f relay, comprising a lens (ACT508-300-A, Thorlabs) and a high-numerical aperture objective (CFI Apo TIRF 100× oil immersion objective, NA = 1.49, Nikon), projected the structured illumination onto the sample. The radius of the annulus on the back focal plane of the objective lens was 3.73 mm, corresponding to an excitation NA of 1.4.

Fluorescence emission was separated from excitation by a multiband dichroic mirror (ZT405/488/561/640-phase R, Chroma) and spectrally filtered using bandpass emission filters centered at 525 nm (ET525/50m), 600 nm (ET600/50m), and 700 nm (ET700/50m) for 488 nm, 561 nm, and 638 nm excitation, respectively.

### System synchronization and acquisition

The timing diagram of the SIM-fdFLIM system is shown in **Supplementary Fig. 1**. Laser modulation was driven by the timing output of the camera operating in external-trigger mode, while the SLM controlled the display of structured illumination patterns. Each SLM diffraction state was synchronized with eight phase-shifted camera exposures, and the nine required SIM patterns were displayed sequentially, generating 8 × 9 raw frames per SIM-fdFLIM cycle.

The overall acquisition rate was primarily limited by the exposure time. With a 1 ms exposure per tap (alternating operation; 72 frames per super-resolved reconstruction), the exposure-time-limited maximum SIM-fdFLIM rates were estimated to be ∼13.9 Hz with eight phase shifts and ∼27.7 Hz with four phase shifts. This estimation does not include additional overhead from camera readout, data transfer, or synchronization delays. For live-cell mitochondrial imaging, longer exposures were used to ensure sufficient signal for SIM reconstruction, resulting in a maximum effective imaging rate of approximately 2.8 Hz with 56 × 56 μm^2^ FOV (**Supplementary Video 4**).

### Cell culture

COS-7 cells (ATCC, CRL-1651) were cultured in Dulbecco’s Modified Eagle’s Medium (DMEM; Gibco, C11995500BT) supplemented with 10% fetal bovine serum (FBS; MeiJing, MJ04001) and 1% penicillin– streptomycin (Gibco, 15140122) at 37 °C in a humidified incubator with 5% CO₂. Cells were seeded at ∼30% confluency onto glass-bottom dishes (Cellvis, D35-20-1.5-N) and grown for 24–36 h before fixation or live-cell imaging.

### FLIM imaging conditions

#### Immunofluorescence imaging (**Fig. 2**)

Cells were rinsed with pre-warmed PBS (HyClone, SH30256.01) before fixation. For microtubule labeling, cells were pre-permeabilized with 0.01% Triton X-100 (Beyotime, ST795) for 5 s at room temperature and fixed with precooled methanol (TGREAG, 105100) at −20 °C for 3 min, followed by three washes with PBS (5 min each). Cells were then permeabilized with 0.2% Triton X-100 for 10 min, washed three times with PBS, and blocked with 10% goat serum (Gibco, 16210064) for 1 h. Primary antibodies against α-tubulin and β-tubulin (Sigma‒Aldrich, T5168 and T5293, respectively; 1:1,000) diluted in 5% goat serum were incubated overnight at 4 °C. After three washes with PBS, cells were incubated for 1 h at room temperature with goat anti-mouse IgG H&L secondary antibodies conjugated to Alexa Fluor 488 (Abcam, ab150113, 1:1,000), Alexa Fluor 546 (Invitrogen, A-11030, 1:500), or Alexa Fluor 647 (Abcam, ab150115, 1:500) diluted in 5% goat serum. Cells were washed three times with PBS and maintained in PBS for imaging. Imaging was performed at 37 °C and pH 7.4.

For dual labeling of mitochondrial outer membranes and microtubules, cells were fixed with 3% paraformaldehyde (Electron Microscopy Sciences, 15710) and 0.1% glutaraldehyde (Electron Microscopy Sciences, 16020) for 15 min at room temperature. After three washes with PBS, cells were permeabilized with 0.2% Triton X-100 for 15 min, blocked with 10% goat serum (Gibco, 16210064) for 1 h, and incubated overnight at 4 °C with mouse anti-α-tubulin, mouse anti-β-tubulin, and rabbit anti-Tom20 (Abcam, ab186735, 1:500) antibodies diluted in 5% goat serum. Following three washes with 0.1% Triton X-100 in PBS, cells were incubated for 1 h at room temperature with goat anti-mouse IgG H&L (Alexa Fluor 546) and goat anti-rabbit IgG H&L (Alexa Fluor 555) (Cell Signaling Technology, 4413S, 1:500) secondary antibodies diluted in 5% goat serum. Cells were then washed three times with 0.1% Triton X-100 in PBS and maintained in PBS for imaging. Imaging was performed at 37 °C and pH 7.4. **Mitochondrial membrane potential imaging** (**Fig. 3**): Cells were incubated with culture medium supplemented with 250 nM PK Mito Red (Genvivo, PKMR-1) at 37 °C for 15 min, followed by washing with pre-warmed DMEM. Images were acquired immediately before (0 min) and 2 min after addition of 10 μM FCCP (Proteintech, CM00253) or 10 μM oligomycin (Proteintech, CM14708). Imaging was performed at 37 °C and pH 7.4 in culture medium.

#### Nile Red membrane staining (**Fig. 4**)

Cells were incubated with culture medium containing 2 μM Nile Red (Invitrogen, N1142) at 37 °C for 45 min and imaged directly. For lipid perturbation experiments, cells were treated with 200 μM oleic acid (OA; Sigma-Aldrich, 364525) overnight, 5 nM paclitaxel (PTX; MedChemExpress, HY-B0015) for 6 h or 4 μM DGAT1 inhibitor (Selleck Chemicals, S7192) together with 4 μM DGAT2 inhibitor (Sigma-Aldrich, PZ0233) overnight prior to staining. Imaging was performed at 37 °C and pH 7.4 in culture medium.

### Image processing and statistical analysis

All image processing was performed using custom Python scripts. Raw camera-acquired SIM-fdFLIM data were corrected for background and preprocessed using the DC-combined lifetime calculation to improve SNR. Super-resolution reconstruction was performed using the Hessian-SIM algorithm^28^. For baseline comparisons, Raw SIM-fdFLIM (termed Raw SIM) denotes SIM reconstruction of the DC-combined SIN and COS channels followed by direct ratio-based lifetime calculation without RIVE. Where indicated, Wiener-SIM was used for this non-RIVE baseline; denoised SIM-fdFLIM denotes Hessian-SIM reconstruction followed by RIVE. Following reconstruction, CC- and TV-regularized ratio-domain denoising (RIVE) was applied to obtain refined lifetime and intensity maps. RIVE was performed with fixed hyperparameters for all datasets (**Supplementary Note 2**).

Structural continuity of SIM reconstructions was quantified using the structure variance metric^28^. The variance of intensity distribution along segmented structures was calculated for each region of interest (ROI), with lower variance indicating smoother and more continuous reconstructed structures. Lifetime-intensity composite images were visualized with intensity encoded as brightness and lifetime encoded as hue. For fixed-cell experiments, line profiles, FWHM measurements, and spatial resolution analyses were performed using ImageJ and MATLAB. For live-cell imaging experiments (Nile Red and PKMR), ROIs were manually selected from intensity images for extraction of temporal intensity and lifetime traces, and mitochondrial dynamics were analyzed by tracking individual organelles across frames. For organelle contact analysis in Nile Red experiments, circular ROIs with a diameter of 112 nm, corresponding to a four-pixel measurement window that approximates the measured lateral resolution of SIM-fdFLIM, were positioned on both sides of adjacent organelles. Overlapping regions and out-of-focus areas were excluded by comparison across at least 10 consecutive frames. For WF data, ROIs were centered at the same positions as in SIM images, while the ROI diameter was increased to 224 nm to approximate diffraction-limited sampling.

Statistical analyses were performed using Origin and MATLAB. Sample sizes, statistical tests, and significance criteria are reported in the corresponding figure legends.

### Reproducibility

Fluorescent dyes were used as references to calibrate the system across different wavelength channels. For 488 nm, 561 nm and 638 nm excitation, antibodies conjugated to Alexa Fluor 488, Alexa Fluor 546 or Alexa Fluor 647 in PBS (1:400 dilution) were used as reference samples respectively, with lifetimes specified by the manufacturer (Thermo Fisher Scientific). Reference measurements were performed every 24 h at 22 °C to account for potential system drift during subsequent sample calibration. All lifetime measurements were calibrated using reference standards acquired with identical modulation frequency, optics, and detector settings. All intensity and lifetime reconstructions were performed using identical parameters, as detailed in the **Supplementary Note**. Hardware parameters, acquisition conditions, and analysis metadata are reported to support reproducibility.

## Data availability

Several example datasets are available via Figshare at https://figshare.com/s/352482f7682bce50cfec. All of the data that support the findings of this study are available from the corresponding author upon reasonable request. Source data are provided with this paper.

## Code availability

The custom code for SIM-fdFLIM reconstruction is available via GitHub at https://github.com/lcy214/SF.git.

## Acknowledgements

This work was primarily supported by the National Key Research and Development Program of China (grant no. 2022YFC3400600 to L.C.) and the National Natural Science Foundation of China (grant nos. 82525105 and 32227802 to L.C.). J.F. acknowledges support from the National Natural Science Foundation of China (grant no. 62576055) and the Natural Science Foundation of Chongqing (grant no. CSTB2025YITP-QCRCX0010). This work was also supported by the Biomedical Computing Platform of the National Biomedical Imaging Center, Peking University. X.H. acknowledges support from the National Key Research and Development Program of China (grant no. 2022YFF0712500), the National Natural Science Foundation of China (grant nos. 32541031, 32322049, and 32350009), the Beijing Nova Programme Interdisciplinary Cooperation Project (grant no. 20240484549), the Beijing Natural Science Foundation (grant no. Z240009), and the Clinical Medicine Plus X-Young Scholars Project of Peking University.

## Author contributions

L.C., J.F., X.H., and B.L. supervised the project. L.C., J.F., and X.H. conceived and initiated the research. C.L. designed the optical layout, built the imaging system, developed the algorithms, and implemented the corresponding software under the guidance of L.C., J.F., and X.H. L.L. prepared the samples and performed the experiments under the guidance of L.C. and X.H. C.L. and L.L. analyzed the data and prepared the figures. L.C., C.L., J.F., L.L., and X.H. wrote the manuscript with input from all authors. All authors participated in data interpretation and scientific discussions.

## References

1. Gustafsson, M. G. L. Surpassing the lateral resolution limit by a factor of two using structured illumination microscopy. J. Microsc. 198, 82–87 (2000).

2. Hell, S. W. & Wichmann, J. Breaking the diffraction resolution limit by stimulated emission: stimulated-emission-depletion fluorescence microscopy. Opt. Lett. 19, 780–782 (1994).

3. Betzig, E. et al. Imaging Intracellular Fluorescent Proteins at Nanometer Resolution. Science 313, 1642–1645 (2006).

4. Rust, M. J., Bates, M. & Zhuang, X. Sub-diffraction-limit imaging by stochastic optical reconstruction microscopy (STORM). Nat. Methods 3, 793–796 (2006).

5. Huang, B., Bates, M. & Zhuang, X. Super-Resolution Fluorescence Microscopy. Annu. Rev. Biochem. 78, 993– 1016 (2009).

6. Zhou, J., del Rosal, B., Jaque, D., Uchiyama, S. & Jin, D. Advances and challenges for fluorescence nanothermometry. Nat. Methods 17, 967–980 (2020).

7. Colom, A. et al. A fluorescent membrane tension probe. Nat. Chem. 10, 1118–1125 (2018).

8. Evans, P. R., Yan, L. & Yasuda, R. Imaging Neuronal Signal Transduction Using Multiphoton FRET-FLIM. in Multiphoton Microscopy (ed. Hartveit, E.) 111–130 (Springer, New York, NY, 2019). doi:10.1007/978-1-4939-9702-2_6.

9. Datta, R., Heaster, T. M., Sharick, J. T., Gillette, A. A. & Skala, M. C. Fluorescence lifetime imaging microscopy: fundamentals and advances in instrumentation, analysis, and applications. J. Biomed. Opt. 25, 071203 (2020).

10. Wang, X. F., Periasamy, A., Herman, B. & Coleman, D. M. Fluorescence Lifetime Imaging Microscopy (FLIM): Instrumentation and Applications. Crit. Rev. Anal. Chem. 23, 369–395 (1992).

11. Auksorius, E. et al. Stimulated emission depletion microscopy with a supercontinuum source and fluorescence lifetime imaging. Opt. Lett. 33, 113–115 (2008).

12. Castello, M. et al. A robust and versatile platform for image scanning microscopy enabling super-resolution FLIM. Nat. Methods 16, 175–178 (2019).

13. Radmacher, N. et al. Doubling the resolution of fluorescence-lifetime single-molecule localization microscopy with image scanning microscopy. Nat. Photonics 18, 1059–1066 (2024).

14. Thiele, J. C. et al. Confocal Fluorescence-Lifetime Single-Molecule Localization Microscopy. ACS Nano 14, 14190–14200 (2020).

15. Shcheslavskiy, V. I. et al. Fluorescence Lifetime Imaging Techniques—A Review on Principles, Applications and Clinical Relevance. J. Biophotonics 18, e202400450 (2025).

16. Gustafsson, M. G. L. et al. Three-Dimensional Resolution Doubling in Wide-Field Fluorescence Microscopy by Structured Illumination. Biophys. J. 94, 4957–4970 (2008).

17. Kner, P., Chhun, B. B., Griffis, E. R., Winoto, L. & Gustafsson, M. G. L. Super-resolution video microscopy of live cells by structured illumination. Nat. Methods 6, 339–342 (2009).

18. Ströhl, F. & Kaminski, C. F. Frontiers in structured illumination microscopy. Optica 3, 667 (2016).

19. Marchand, R. et al. Super-resolution structured illumination microscopy of cellular dynamics with fluorescence lifetime contrast. Optica 13, 1065 (2026).

20. Gadella, T. W. J., Jovin, T. M. & Clegg, R. M. Fluorescence lifetime imaging microscopy (FLIM): Spatial resolution of microstructures on the nanosecond time scale. Biophys. Chem. 48, 221–239 (1993).

21. Chen, H., Holst, G. & Gratton, E. Modulated CMOS camera for fluorescence lifetime microscopy. Microsc. Res. Tech. 78, 1075–1081 (2015).

22. Smith, C. S. et al. Structured illumination microscopy with noise-controlled image reconstructions. Nat. Methods 18, 821–828 (2021).

23. Chen, X. et al. Superresolution structured illumination microscopy reconstruction algorithms: a review. Light Sci. Appl. 12, 172 (2023).

24. Chang, C.-W. & Mycek, M.-A. Precise fluorophore lifetime mapping in live-cell, multi-photon excitation microscopy. Opt. Express 18, 8688–8696 (2010).

25. Spring, B. Q. & Clegg, R. M. Image analysis for denoising full-field frequency-domain fluorescence lifetime images. J. Microsc. 235, 221–237 (2009).

26. Franke, R. & Holst, G. A. Frequency-domain fluorescence lifetime imaging system (pco.flim) based on an in-pixel dual tap control CMOS image sensor. in Imaging, Manipulation, and Analysis of Biomolecules, Cells, and Tissues XIII vol. 9328 241–259 (SPIE, 2015).

27. Raspe, M. et al. siFLIM: single-image frequency-domain FLIM provides fast and photon-efficient lifetime data. Nat. Methods 13, 501–504 (2016).

28. Huang, X. et al. Fast, long-term, super-resolution imaging with Hessian structured illumination microscopy. Nat. Biotechnol. 36, 451–459 (2018).

29. Shannon, C. E. A Mathematical Theory of Communication. Bell Syst. Tech. J. 27, 379–423 (1948).

30. Thompson, R. E., Larson, D. R. & Webb, W. W. Precise Nanometer Localization Analysis for Individual Fluorescent Probes. Biophys. J. 82, 2775–2783 (2002).

31. Van Trees, H. L. Detection, Estimation, and Modulation Theory. (New York, Wiley, 1968).

32. Bevington, P. R. Data Reduction and Error Analysis for the Physical Sciences. (Boston : McGraw-Hill, 2003).

33. Müller, M., Mönkemöller, V., Hennig, S., Hübner, W. & Huser, T. Open-source image reconstruction of super-resolution structured illumination microscopy data in ImageJ. Nat. Commun. 7, 10980 (2016).

34. Saimi, D. et al. Imaging mitochondrial membrane potential via concentration-dependent fluorescence lifetime changes. Nat. Commun. 16, 11088 (2025).

35. Wolf, D. M. et al. Individual cristae within the same mitochondrion display different membrane potentials and are functionally independent. EMBO J. 38, e101056 (2019).

36. Mukherjee, S., Raghuraman, H. & Chattopadhyay, A. Membrane localization and dynamics of Nile Red: Effect of cholesterol. Biochim. Biophys. Acta BBA - Biomembr. 1768, 59–66 (2007).

37. Gajo, C. et al. Nile Red Fluorescence: Where’s the Twist? J. Phys. Chem. B 128, 11768–11775 (2024).

38. Greenspan, P., Mayer, E. P. & Fowler, S. D. Nile red: a selective fluorescent stain for intracellular lipid droplets. J. Cell Biol. 100, 965–973 (1985).

39. Zhanghao, K. et al. High-dimensional super-resolution imaging reveals heterogeneity and dynamics of subcellular lipid membranes. Nat. Commun. 11, 5890 (2020).

40. Levitt, J. A., Chung, P.-H. & Suhling, K. Spectrally resolved fluorescence lifetime imaging of Nile red for measurements of intracellular polarity. J. Biomed. Opt. 20, 096002 (2015).

41. Song, Y. S., Won, Y. J. & Kim, D. Y. Time-lapse in situ fluorescence lifetime imaging of lipid droplets in differentiating 3T3-L1 preadipocytes with Nile Red. Curr. Appl. Phys. 15, 1634–1640 (2015).

42. Bianchetti, G., et al. Investigation of the Membrane Fluidity Regulation of Fatty Acid Intracellular Distribution by Fluorescence Lifetime Imaging of Novel Polarity Sensitive Fluorescent Derivatives. Int. J. Mol. Sci. 22, 3106 (2021).

43. Zeug, A., Woehler, A., Neher, E. & Ponimaskin, E. G. Quantitative Intensity-Based FRET Approaches—A Comparative Snapshot. Biophys. J. 103, 1821–1827 (2012).

44. Kao, J. P. Y., Li, G. & Auston, D. A. Practical aspects of measuring intracellular calcium signals with fluorescent indicators. Methods Cell Biol. 99, 113–152 (2010).

45. Dean, W. F. & Mattheyses, A. L. Illuminating cellular architecture and dynamics with fluorescence polarization microscopy. J. Cell Sci. 137, jcs261947 (2024).

46. Fogarty, M. P. & Warner, I. M. Ratio method for fluorescence spectral deconvolution. Anal. Chem. 53, 259–265 (1981).

47. Moon, S. et al. Spectrally Resolved, Functional Super-Resolution Microscopy Reveals Nanoscale Compositional Heterogeneity in Live-Cell Membranes. J. Am. Chem. Soc. 139, 10944–10947 (2017).

48. Pott, E. D., Yang, M., Batey, J. E., Embree, J. & Dong, B. Fast Hyperspectral and Super-Resolved Mapping of Lipid Membrane Polarity with Single-Molecule Sensitivity. Chem. Biomed. Imaging 4, 584–589 (2026).

