## Supplementary material for "Ratio-preserving super-resolution FLIM for quantitative live-cell lifetime imaging"

**Supplementary Note 1 – Mathematical description of SIM-fdFLIM**

In an fdFLIM system, the illumination is periodically modulated, which induces a fluorescence signal that fluctuates at the same frequency. For sinusoidally modulated illumination, the fluorescence emission can be written as:

$$e(t) = A[1 + m\sin(2\pi f \cdot t + \varphi_\tau + \varphi_r)] \tag{1}$$

where $A$ denotes the fluorescence amplitude determined by the sample, $m$ corresponds to the modulation depth of the fluorescence signal, $f$ is the modulation frequency of the excitation and fluorescence emission signals, $t$ represents time, $\varphi_\tau$ is the phase difference between illumination and fluorescence and $\varphi_r$ is the system-induced phase offset. Each harmonic (order) component $a_k$ of the fluorescence emission $e(t)$ can be obtained by Fourier transform:

$$a_k = \frac{1}{T}\int_0^T e(t)\exp[-i(k \cdot 2\pi ft + \varphi_r)]dt = G_k + iS_k \tag{2}$$

where the number of orders is $k$, the period of signal is $T$, $G_k$ and $S_k$ are the $k^{th}$ order cosine and sine components, respectively. $\varphi_r$ is employed in the transformation to eliminate the system induced phase, which can be obtained from the reference fluorophore with known and stable lifetime[1,2]. The first-order components are related to the modulation properties of the fluorescence signal, whereas the zero-order component is the constant (DC) term:

$$G_0 = A;\ S_0 = 0;\ G_1 = \frac{Am}{2}\sin\varphi_\tau;\ S_1 = \frac{Am}{2}\cos\varphi_\tau \tag{3}$$

Therefore, the fluorescence lifetime $\tau$ of the fluorophore can be calculated from the ratio of the first-order components:

$$\tau = \frac{\tan\varphi_\tau}{2\pi f} = \frac{1}{2\pi f}\frac{G_1}{S_1} \tag{4}$$

In the commercially available FLIM camera used in this work, the periodically modulated fluorescence signal is split into two captures with an initial phase offset of $\pi$. By changing the initial phase of this pair, an even number of images with equally spaced initial phases can be acquired (**Fig. 1**). Discrete Fourier analysis of the temporally sampled signal yields the DC component and the first-harmonic sine and cosine components:

$$G_0 = \frac{1}{q}\sum_{n=1}^{q} Y(n);\ G_1 = \sum_{n=1}^{q} Y(n)\cdot\cos(n\Delta\varphi + \varphi_r);\ S_1 = \sum_{n=1}^{q} Y(n)\cdot\sin(n\Delta\varphi + \varphi_r) \tag{5}$$

Here, $q$ is the number of initial phases, $\Delta\varphi$ is the interval between the initial phases of successive captures (a constant, $\pi/4$ in our case), and $Y(n)$ is the measured fluorescence signal for each initial phase, recorded with the same exposure time and phase interval $\pi$. The measured intensity signal can be expressed as:

$$Y(n) = \int_{\varphi_n}^{\varphi_n+\pi} I(\vec{p},t) \cdot e(t) \otimes \mathrm{PSF}\ dt \tag{6}$$

where $\varphi_n$ is the initial phase of the capture, $\vec{p}$ is the position vector on the x-y plane, $t$ represents time, $I(\vec{p},t)$ is the structured illumination intensity, $\otimes$ means convolution and PSF is the point spread function of the optical system. For lifetime measurement and SIM reconstruction, the structured illumination is both spatially and temporally modulated in a sinusoidal form:

$$I(\vec{p},t) = A_t[1 + m_t \exp(ik_t t + \varphi_\tau)] \cdot A_{\vec{p}}[1 + m_{\vec{p}} \exp(i\vec{k}_{\vec{p}} \cdot \vec{p} + \varphi_{\vec{p}})] \tag{7}$$

Here, $A_t$ and $A_{\vec{p}}$ are the amplitudes, $m_t$ and $m_{\vec{p}}$ are the modulation depths, $k_t = 2\pi f$ and $k_{\vec{p}}$ are the spatial frequency vectors, $\varphi_\tau$ and $\varphi_{\vec{p}}$ are the initial phases of the temporal and spatial modulations, respectively. The phase $\varphi_\tau$ is related to fluorescence lifetime. Since the three components $G_0$, $G_1$ and $S_1$ are linear combinations of the same spatially modulated signal $Y(n)$, the illumination modulation depth in the x-y plane is preserved. Therefore, these three components can be reconstructed into super-resolution SIM (SR-SIM) images respectively. Consequently, taking the ratio between the two first-order components in the reconstructed images should lead to the SR lifetime distribution (**Eq.(5)**).

However, the first-order components $G_1$ and $S_1$ obtained directly from Eq.(5) often have relatively low signal-to-noise ratio (SNR), and their values can even become negative because of additional system-induced phase shifts. To enhance SNR and ensure high-quality SIM reconstruction, we incorporate the DC component into the lifetime calculation. We define $X_{\cos}$ and $X_{\sin}$ as the following:

$$X_{\cos} = M \cdot G_0 + G_1 \tag{8}$$
$$X_{\sin} = M' \cdot G_0 + S_1 \tag{9}$$

where $M$ and $M'$ are constant coefficients. In practice, $X_{\cos}$ and $X_{\sin}$ can be calculated according to **Eq.(5)**:

$$X_{\cos} = \frac{M}{q}\sum_{n=1}^{q} Y(n) + \sum_{n=1}^{q} Y(\mathrm{n}) \cdot \cos(n\Delta\varphi + \varphi_r) \tag{10}$$

$$X_{\sin} = \frac{M'}{q}\sum_{n=1}^{q} Y(n) + \sum_{n=1}^{q} Y(\mathrm{n}) \cdot \sin(n\Delta\varphi + \varphi_r) \tag{11}$$

where $X_{\cos}$ and $X_{\sin}$ both own high SNR satisfying the need of high-quality SIM reconstruction. The widefield intensity images and corresponding SIM results from the traditional and DC-combined methods are shown in **Supplementary Fig. 2c**.

When a fluorophore exhibits single-exponential decay, the lifetime obtained from modulation depth is identical to that obtained from phase shift, and the emission modulation depth is $m = m_t \cos\varphi_\tau$. After SIM reconstruction, the ratio of the two SR results can be written according to **Eq.(3)**:

$$F = \frac{X_{re-cos}}{X_{re-sin}} = \frac{M \tan^2 \varphi_\tau + M + 0.5 m_t \tan \varphi_\tau}{M' \tan^2 \varphi_\tau + M' + 0.5 m_t} \quad (12)$$

where $X_{\text{re-cos}}$ and $X_{\text{re-sin}}$ are the SIM reconstructed results of $X_{\cos}$ and $X_{\sin}$, $\tan\varphi_\tau \in (0, \infty)$ and $\varphi_\tau \in (0, \pi/2)$. Widefield mode can be written as $F = X_{a\text{-cos}} / X_{a\text{-sin}}$, where $X_{a\text{-cos}}$ and $X_{a\text{-sin}}$ are approximated by averaging $X_{\cos}$ and $X_{\sin}$ across different spatial patterns. In the possible range of $m_t \in [0.5, 1]$, the function $F$ is almost monotonically increasing when $M'$ does not exceed $M$. Functions with different $M$ and $M'$ are illustrated in **Supplementary Fig. 2b**, all of which preserve a monotonic mapping to lifetime within $[0, 25]$ ns under a 20 MHz modulation frequency. The slope of the curve is influenced by $m_t$ in a monotonic manner (**Supplementary Fig. 2b**), which is decided by illumination laser and exposure time. $m_t$ for a given imaging condition can be calculated from the reference fluorophore with known lifetime as well. This working range can be further extended by employing a lower modulation frequency. Therefore, the mapping from $F$ to $\varphi_\tau$ can be used to infer $\varphi_\tau$ and lifetime in a consistent and physically meaningful manner.

However, although the stabilized $F$ metric demonstrates improved robustness to noise, the incorporation of the DC component reduces its sensitivity to changes in lifetime, as reflected by the slope curves in **Supplementary Fig. 2b**. To maintain sensitivity comparable to the conventional frequency-domain method, we select $M = M' = 1/8$ in our system for the detection of lifetimes below 10 ns. Notably, in background regions with weak fluorescence signals, the modulation depth approaches zero, causing $F$ to converge to a constant value $M'/M$. This property enables background suppression directly in the lifetime image. When the target lifetime is well separated from this limiting value (~7.9 ns in our case), a simple threshold can be applied to distinguish signal from background (**Supplementary Fig. 3**). In practice, even more exaggerated values can be observed in background regions due to random noise. For the $M = M' = 1/8$ curve, the slope decreases markedly beyond $F = M'/M = 1$, such that noise-induced perturbations in $F$ can lead to disproportionately large lifetime estimates. Overall, compared with the conventional approach of $G_1 / S_1$, the DC-combined method improves the SNR, contributing to both more accurate lifetime estimation and higher-quality SIM reconstructions (**Supplementary Fig. 2**).

### Supplementary Note 2 – The denoising SIM-fdFLIM reconstruction algorithm

The $X_{\cos}$ and $X_{\sin}$ of **Eq.(8)** and **Eq.(9)** can be reconstructed directly to obtain the SIM intensity images. However, calculating SR lifetime requires division between reconstructed channels, which amplifies noise and produces artifacts in lifetime images (**Supplementary Fig. 3**). To address this problem, we propose a two-step denoising SIM-fdFLIM reconstruction algorithm that yields accurate SR lifetime distributions.

First, to suppress artifacts introduced by sample motion and noise in intensity images, we use Hessian-SIM algorithm to reconstruct the SIM images of $X_{cos}$ and $X_{sin}$ [3]. This step is essential in reconstructing SIM images from raw data of low SNR, which are inevitable during the acquisition of different modulation depths of the fluorescence signals.

Direct calculation of lifetime from the ratio of $X_{re-cos}$ and $X_{re-sin}$ (Hessian-SIM reconstructed $X_{cos}$ and $X_{sin}$) is possible. However, despite the denoising of intensity images by the Hessian regularization, direct division suffers from distortion induced by minor mismatched paired images and artifacts caused by noise amplification. Therefore, we propose ratio-domain integrated variational estimation (RIVE), which reformulates the division operation as a multiplicative iterative model and incorporates multiple regularization terms to stabilize lifetime recovery.

Because of the finite system resolution and randomly distributed noise, a slight sub-pixel mismatch can exist between the $X_{re-cos}$ and $X_{re-sin}$ channels, even though their overall intensity structures are nearly identical. This mismatch has a negligible effect on the intensity images ($X_{re}$) but can severely distort the ratio image ($k$), manifesting as strong spatial phase misplacement between $X_{re}$ and $k$ (**Supplementary Fig. 5**). To mitigate this, we introduced a cross-correlation (CC) penalty term to maximize the similarity between the structural channels and the ratio channel. However, prior to CC, we needed to normalize $X_{re-cos}$ and $X_{re-sin}$ to better incorporate information from the structural channel to reciprocally constrain the $k$ channel. As biological samples typically show regions of signals of high SNR and background of low SNR, we divided the $X_{re-cos}$ and $X_{re-sin}$ images into local blocks and performed signal-aware normalization within each block, which improved the overall lifetime reconstruction. In fact, the application of a global CC penalty without block partition led to failure in resolving lifetime difference of weakly labeled structure from the background in the final reconstruction, whereas the $k$ image with CC penalty applied with incorrectly over-partitioned blocks exhibited amplified artifacts at regions of low SNR (**Supplementary Fig. 4**). In contrast, a partitioned normalization step with 4×4 partition block produced lifetime reconstruction of good balance between sensitivity and artifacts generation. For our data, a 4×4 partition for the field of view of 56×56 $\mu m^2$ works well. Overall, such a signal-aware, partitioned normalization step enhances the sensitivity of the subsequent CC penalty terms between the normalized $X_{re-cos}$ and $X_{re-sin}$ with the $k$, enabling maximal and robust subpixel alignments between the intensity and the ratio images.

Second, we add a total variation (TV) penalty to further regularize the ratio image, since noise may persist even after mismatch correction with the CC penalty. Other regularization terms, such as Hessian penalties, can also provide denoising and smoothing. However, TV is particularly suitable for images with piecewise-uniform regions and sharp edges, which matches the typical structure of lifetime maps. Therefore, we choose TV, rather than Hessian, to regularize the ratio term in our algorithm. Finally, because $X_{\text{re-cos}}$ and $X_{\text{re-sin}}$, and $k$ channels may have different noise distributions, we construct the complete optimization function as a symmetric loss function to prevent convergence collapse (**Eq.(13)**):

$$\begin{aligned}\min_{k_1,k_2} \| X_{\text{re-cos}} - k_1 \cdot X_{\text{re-sin}} \|_2^2 + \| X_{\text{re-sin}} - k_2 \cdot X_{\text{re-cos}} \|_2^2 + \mu_1 \| k_1 \|_2^2 + \mu_2 \| k_2 \|_2^2 \\ + \lambda_1 \cdot \mathrm{CC}(X_{\text{re-cos}}, k_1) + \lambda_2 \cdot \mathrm{CC}(X_{\text{re-sin}}, k_2) + \alpha_1 \cdot \mathrm{TV}(k_1) + \alpha_2 \cdot \mathrm{TV}(k_2)\end{aligned} \tag{13}$$

where $k_1$ and $k_2$ are the ratio and inverse ratio of $X_{\text{re-cos}}$ and $X_{\text{re-sin}}$ respectively, $\mathrm{CC}(X,k)$ is the cross-correlation regularization term, $\mathrm{TV}(k)$ is the total variation penalty term. $\mu_1 = 0.08$ and $\mu_2 = 0.072$ are the fidelity term parameters, which restrain the $k$ values from distortion. $\lambda_1 = \lambda_2 = 0.4$, $\alpha_1 = \alpha_2 = 0.05$ are the penalty term parameters. $\mathrm{CC}(X,k)$ and $\mathrm{TV}(k)$ can be obtained as:

$$\mathrm{CC}(X,k) = -\sum_{\vec{p}} X(\vec{p}) \cdot k(\vec{p}) \tag{14}$$

$$\mathrm{TV}(k) = \sum_{\vec{p}=(x,y)} \sqrt{\left(\frac{\partial k}{\partial x}\right)^2 + \left(\frac{\partial k}{\partial y}\right)^2} \tag{15}$$

We minimize this loss function iteratively using gradient descent, with the step sizes determined adaptively by the Adam optimizer[4].

Having completed the full processing pipeline, we evaluated the role of the CC and TV regularization steps in improving the fidelity of ratio images. The effectiveness of mismatch correction is demonstrated using AF488-stained microtubules in **Supplementary Fig. 5**. When various regularization strategies are applied to the same dataset, the centroid distance between the two PSFs is reduced from 0.034 to 0.002 pixels (0.952 to 0.056 nm, 28 nm per pixel) by the combined CC+TV approach. CC alone yields inferior performance due to noise-induced fluctuations, whereas TV shows an apparent alignment effect. However, this behavior of TV results from enforcing a flattened ratio rather than correcting the misalignment. As a consequence, TV alone produces mismatched channels and erroneous lifetime estimates. Hessian regularization, on the other hand, is unsuitable for ratio images because it excessively smooths edges. The respective effects of CC and TV are further illustrated in **Supplementary Fig. 5-6**. CC preserves spatial fluctuations and prevents over-smoothing by TV. The combined CC+TV scheme achieves the best balance, maintaining the highest spatial resolution of 103.5 nm. Meanwhile, the contribution of TV effectively denoises the ratio, yielding a standard deviation comparable to widefield FLIM. The lifetime accuracy and precision achieved by the CC+TV approach are demonstrated in **Supplementary Fig. 7** using single-labeled fixed samples.

Besides, CC+TV regularization is particularly effective at suppressing background noise. Because background noise is randomly distributed and of low absolute intensity, the cross-correlation value is maximized when the ratio becomes a constant relative to regions containing true fluorescence signal. According to the DC-combined FLIM model, the constant is known and can be filtered by a value threshold in most cases (**Supplementary Note 1**). As a result, background fluctuations are suppressed without requiring an explicit structural mask to separate foreground from background (**Supplementary Fig. 3**); threshold masking was applied only for visualization or for defining the quantitative lifetime range. Moreover, CC+TV preserves meaningful inter-channel variations and prevents over-smoothing of the lifetime map. This behavior can be understood from two representative scenarios:

1. **Sub-PSF fluctuations:** For small-scale fluctuations within the size of a point spread function (PSF), which are dominated by noise, both CC and TV suppress these fluctuations by alignment and smoothing, respectively.
2. **Super-PSF variations:** When the ratio varies over spatial ranges larger than the PSF, TV alone tends to smooth out these genuine differences, whereas CC encourages their preservation because they correspond to true intensity relationships between the channels. The symmetric design also prevents the ratio from biasing toward either intensity channel.

This complementary behavior across spatial scales relative to the PSF prevents loss of fidelity while maintaining high spatial resolution in the ratio image. Simulations further demonstrate the fidelity and stability of RIVE-optimized ratio images under different intensity-lifetime relationships, including regions with different lifetimes but identical intensities and regions with identical lifetimes but different intensities (**Supplementary Fig. 9**). Consequently, even in the presence of smoothing-type regularization, the essential spatial variations in the lifetime map are preserved, enabling accurate discrimination of multiple lifetime populations (**Supplementary Fig. 8**).

### Supplementary Note 3 – Discussion

#### 1. Technical considerations in SIM-fdFLIM reconstruction

SIM-fdFLIM combines spatially structured illumination with temporally modulated excitation, which inherently increases the number of raw frames required for reconstruction. In principle, super-resolution lifetime recovery is feasible as long as the total detected photon budget is sufficient; however, traditional reconstruction methods implicitly assume that each individual frame has adequate SNR. This assumption limits achievable temporal resolution and increases susceptibility to photobleaching when short exposure times are used.

The DC-combined lifetime formulation mitigates this limitation by stabilizing phase-based lifetime estimation under low-photon conditions. Inclusion of the DC component improves robustness of both phase and modulation-depth estimates, particularly when detected phase shifts approach $0.5\pi$, and is essential for maintaining sufficient SNR in both channels for SIM reconstruction. For mixed-lifetime samples, the extracted lifetime represents a weighted intermediate of the contributing components, which is appropriate for heterogeneous cellular environments where sub-pixel mixing is unavoidable.

**2. Ratio-domain regularization and limitations**

The CC+TV-regularized iterative reconstruction refines lifetime estimates directly in the ratio domain, improving numerical stability and suppressing noise amplification inherent to division operations. While this approach effectively mitigates random noise and subpixel misalignment between channels, it cannot fully correct for structurally induced low-SNR effects such as out-of-focus fluorescence. In widefield fdFLIM, out-of-focus contributions persist frame-to-frame and can lead to systematic lifetime deviations. In SIM-fdFLIM, these effects are most apparent for thin membrane structures spanning a depth comparable to or exceeding the axial response of the system, such as the mitochondrial outer membrane.

With an effective focal depth of ~400 nm and mitochondrial thickness approaching ~700 nm, out-of-focus contributions can be amplified during super-resolution reconstruction, leading to mixed lifetime values in intermediate regions. Consequently, intensity images remain an essential reference for identifying in-focus regions where lifetime estimates are reliable. These considerations do not alter the main conclusions, but define the spatial contexts in which lifetime values should be interpreted most cautiously.

**3. Interpretation of lifetime contrast in membrane systems**

Fluorescence lifetime of solvatochromic dyes such as Nile Red reflects a composite of local polarity, hydration, microviscosity, protein crowding, and dye concentration, rather than membrane order alone. As a result, shorter lifetimes in densely packed structures such as mitochondrial cristae can arise from increased non-radiative decay pathways, including self-quenching and protein-lipid interfacial effects. Similarly, asymmetric lifetime signatures at organelle contact sites likely reflect differences in local lipid remodeling and probe sensitivity on each side of the interface rather than symmetric chemical environments.

**4. Generalizability and future directions**

Although demonstrated here for SIM-fdFLIM, both the DC-combined lifetime metric and RIVE are broadly applicable to other functional imaging modalities that rely on ratio calculations, including FRET, spectral unmixing, and ratiometric ion indicators. Future developments incorporating adaptive illumination, improved axial sectioning, or machine-learning-assisted reconstruction may further enhance temporal resolution and extend SIM-fdFLIM to volumetric lifetime imaging.

**Reference**


1. Raspe, M. *et al.* siFLIM: single-image frequency-domain FLIM provides fast and photon-efficient lifetime data. *Nat. Methods* **13**, 501–504 (2016).
2. Chen, H. & Gratton, E. A practical implementation of multifrequency widefield frequency-domain fluorescence lifetime imaging microscopy. *Microsc. Res. Tech.* **76**, 282–289 (2013).
3. Huang, X. *et al.* Fast, long-term, super-resolution imaging with Hessian structured illumination microscopy. *Nat. Biotechnol.* **36**, 451–459 (2018).
4. Kingma, D. P. & Ba, J. Adam: A Method for Stochastic Optimization. Preprint at https://doi.org/10.48550/arXiv.1412.6980 (2017).

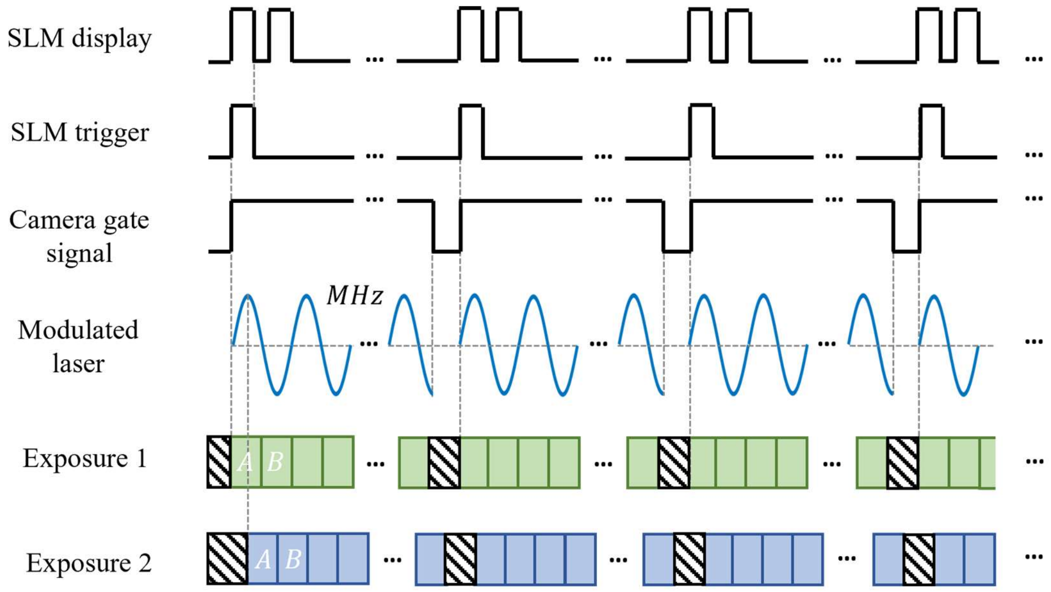


**Supplementary Fig. 1| Timing chart of SIM-fdFLIM system.**

The MHz modulation signal is generated by the dual-tap fdFLIM camera (pco.flim X, PCO), which modulates the laser intensity. The camera gate signal initiates the modulation sequence, shutters the laser during camera readout, and synchronizes the spatial light modulator (SLM). The two charge-collection taps (Tap A and Tap B) of each camera pixel operate alternately, accumulating signals from the same modulation phase period in each cycle. Therefore, each exposure contains two images: two exposures are required for four-sample acquisition, yielding four images, whereas four exposures are required for eight-sample acquisition, yielding eight images. For each lifetime stack, the structured illumination pattern remains unchanged over eight or four phase-shifted exposures; the pattern is then advanced through nine SIM states, yielding 8 × 9 or 4 × 9 raw images per SIM-fdFLIM acquisition.

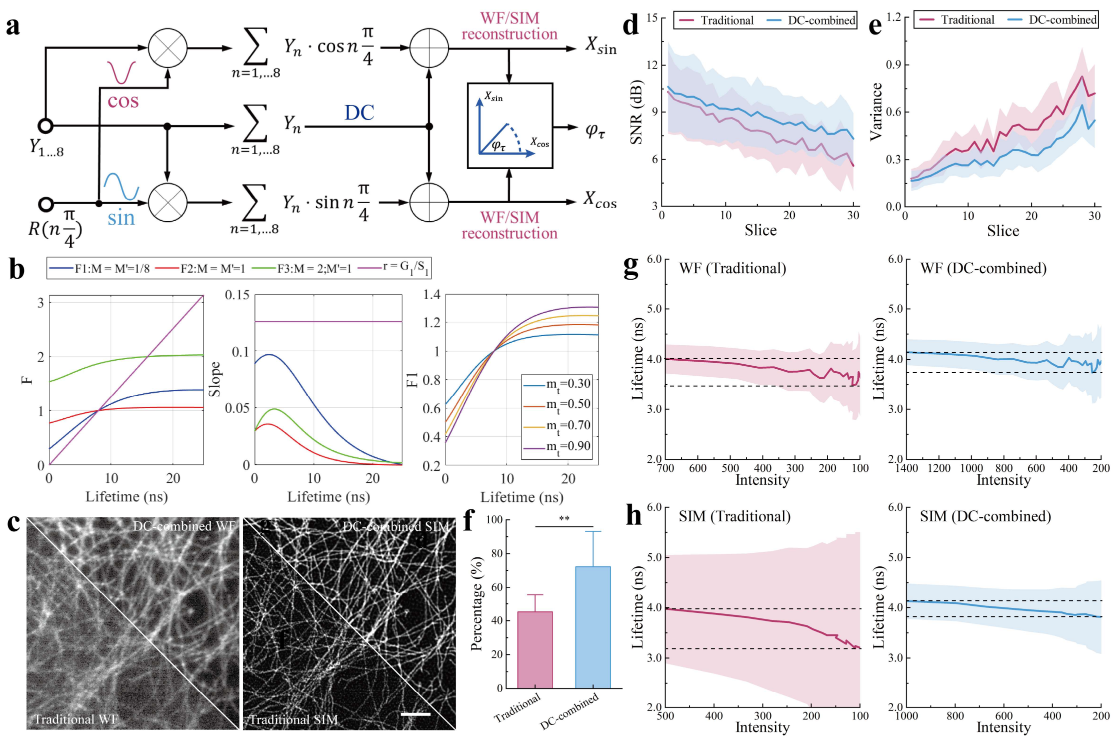


**Supplementary Fig. 2| Direct-current (DC) combined lifetime calculation.**

**a,** Workflow of the DC-combined FLIM calculation. $R$: sine/cosine basis functions for discrete Fourier transform.

**b,** Relationship between F and lifetime with a 0.9 illumination modulation ($m_t$) under different DC scaling factors (left), including r, which corresponds to the traditional calculation. The corresponding slopes computed as ΔF/Δτ are shown in the middle. F1 curves with different $m_t$ are shown on the right.

**c,** Traditional (lower) and DC-combined (upper, using a 1/8 DC component) intensity images in WF (left) and SIM (right).

**d,** WF SNR of an AF488-tubulin sample continuously acquired over 30 frames. Each frame consisted of 72 images acquired with 150 ms exposure time.

**e,** Variance curves of Wiener-reconstructed SIM results without RIVE for the two methods, indicating structural continuity.

**f,** Bar plots showing percentage of traditional and DC-combined SIM reconstructions with variance below 0.5, presented as mean + SD ($n$ = 12 captures per group). **P < 0.01.

**g,** WF lifetime measured over 30 consecutive frames as a function of corresponding sine-channel intensity for the two methods.

**h,** Hessian SIM with RIVE lifetime over 30 consecutive frames as a function of corresponding sine-channel intensity for the two methods.

Scale bar: 1 μm. In **d-e** and **g-h**, shaded regions represent standard deviation (SD), $n \geq 12$ ROIs.

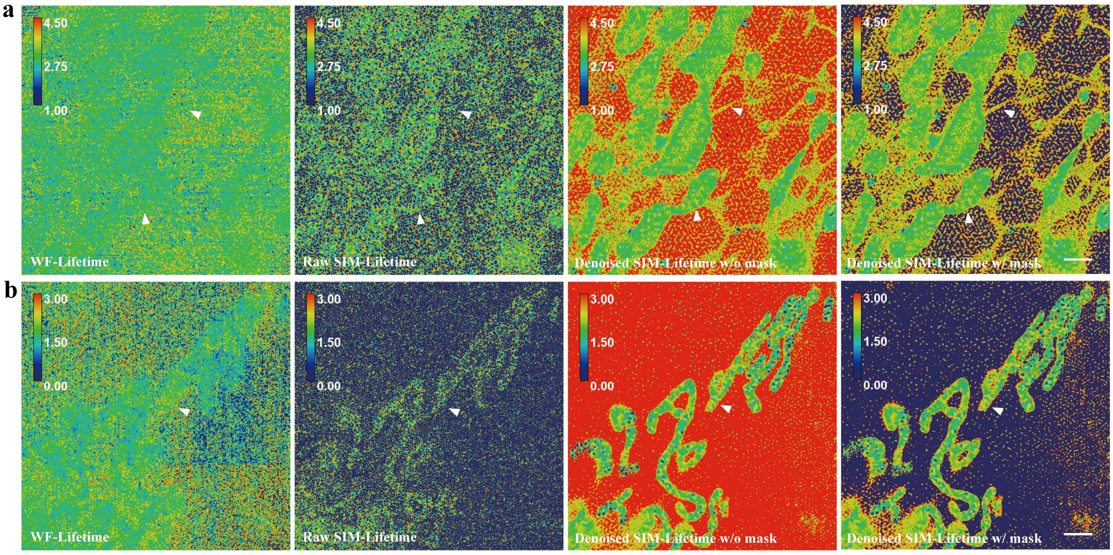


**Supplementary Fig. 3| Live-cell fluorescence lifetime images obtained with different reconstruction modes.**

**a,** Fluorescence lifetime images of Nile Red-stained COS7 cells acquired by WF fdFLIM, raw SIM-fdFLIM (Wiener reconstruction with threshold masking; termed raw SIM), denoised SIM-fdFLIM (Hessian SIM with RIVE; termed denoised SIM) without threshold masking, and denoised SIM-fdFLIM with threshold masking, from left to right.

**b,** Lifetime distribution of COS7 cell stained with PKMR arranged as the same sequence of **a**. Scale bar: 2 μm. For PKMR images, pixels with lifetimes > 3 ns are shown in black for visualization. For Nile Red images, the displayed lifetime range is 1-4.5 ns.

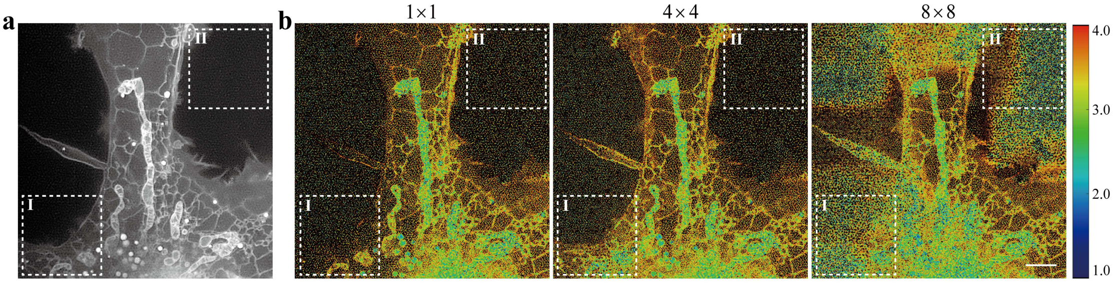


**Supplementary Fig. 4| The effect of partitioning on denoising results.**

**a,** Intensity image of a live COS7 cell labeled with Nile Red.

**b,** Lifetime image of a live COS7 cell labeled with Nile Red. From left to right: RIVE-denoised results obtained using different partitioning schemes (1×1, 4×4, and 8×8). Region I contains weaker signal compared with the centre of the image, leading to signal loss using the 1×1 partition. Region II contains no fluorescence signal. Therefore, excessively fine partitioning produces unstable, low-SNR results in this background-dominated area. Scale bar: 2 μm.

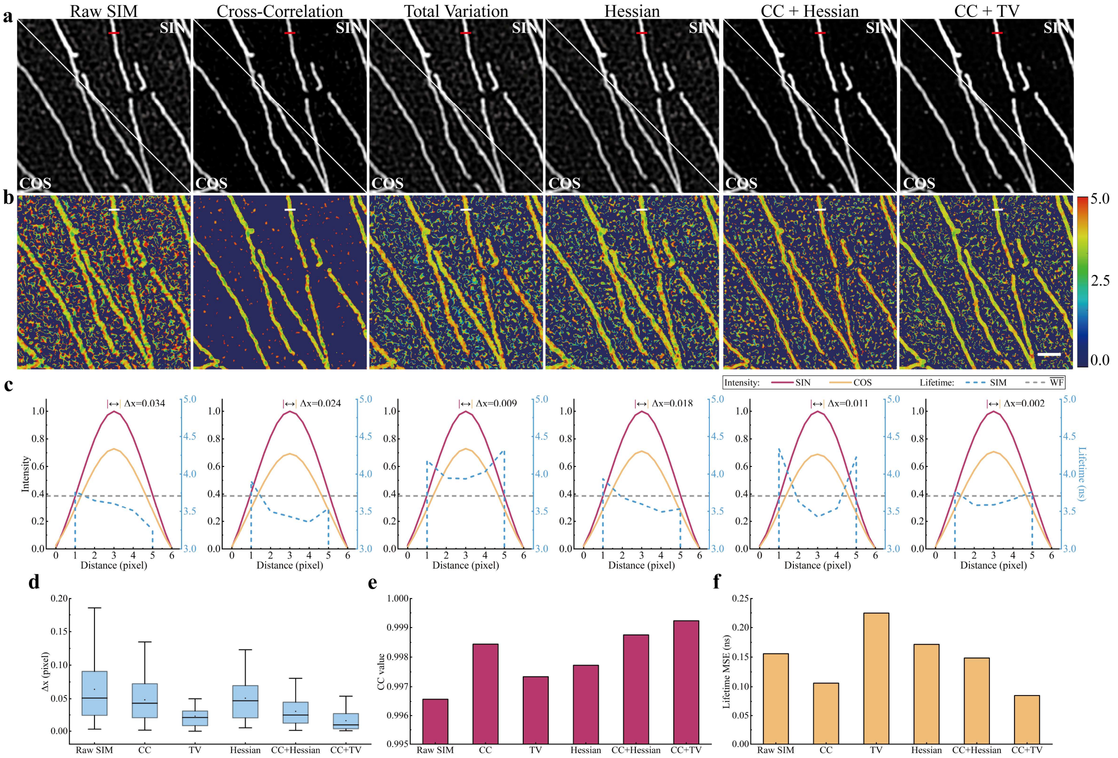


**Supplementary Fig. 5| Mismatch correction performance demonstrated using AF488-stained microtubules in fixed COS-7 cells.**

**a,** Reconstructed SIN and COS intensity images obtained using Wiener-SIM reconstruction only (raw SIM), Hessian-SIM reconstruction with cross-correlation (CC) regularization, Hessian-SIM reconstruction with total variation (TV) regularization, Hessian-SIM reconstruction with Hessian regularization, Hessian-SIM reconstruction with CC + Hessian regularization, and Hessian-SIM reconstruction with CC + TV regularization (from left to right). All SIM results are reconstructed from DC-combined widefield data.

**b,** Lifetime images obtained from the same set of denoising and regularization strategies as in **a**.

**c,** Normalized Gaussian-fitted line profiles of the SIN and COS channels and the corresponding lifetime along the marked line in a and b. Δx denotes the distance between the centroid positions of the SIN and COS profiles. The gray dotted line indicates the WF-fdFLIM lifetime (3.71 ns).

**d,** Box plots showing the centroid distance (Δx) under different regularization strategies. Data are presented as mean (dot), median (center line), interquartile range (box), and 5-95% range (whiskers). $n$ = 24 ROIs per group.

**e,** Bar plots showing the cross-correlation values between the COS and SIN channels in **a** under different regularization strategies.

**f,** Bar plots showing the lifetime mean squared error (MSE) in **b** obtained with different regularizations, using the WF-fdFLIM lifetime map within the same intensity mask as the reference. Scale bar: 500 nm.

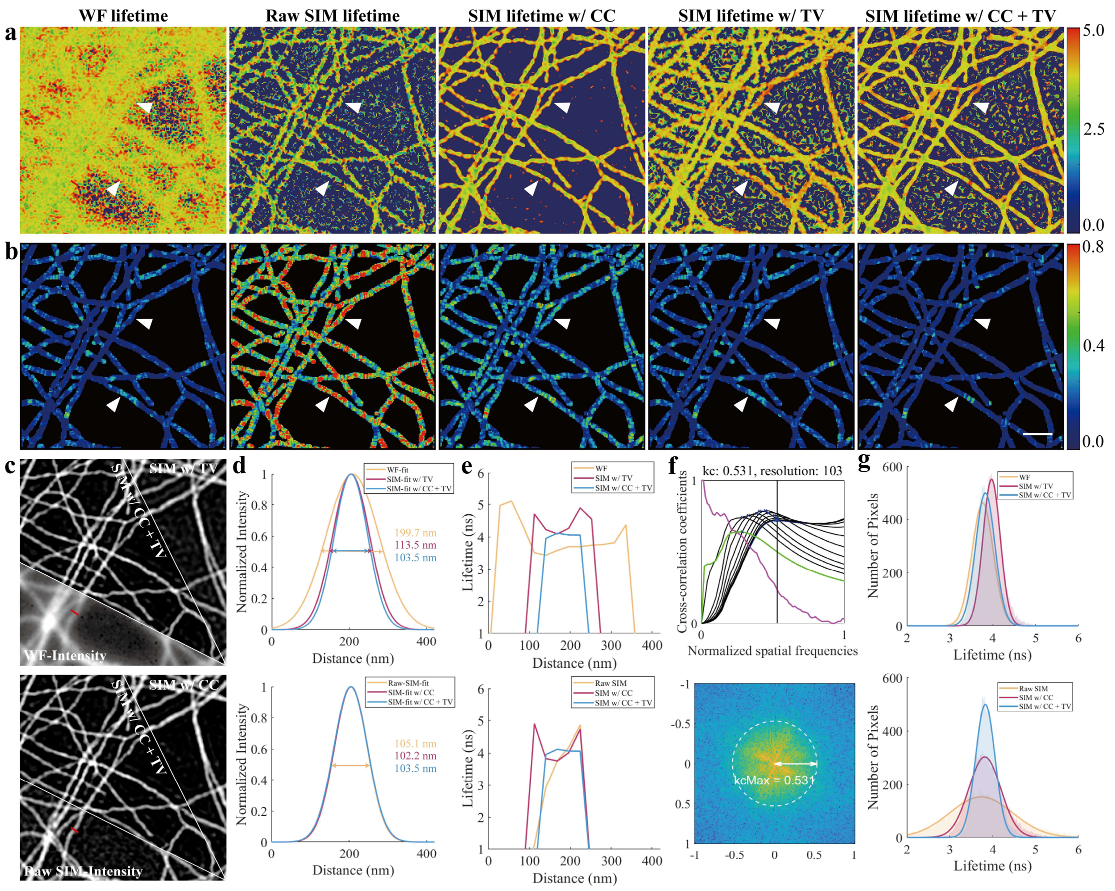


**Supplementary Fig. 6| Resolution enhancement by CC and denoising capability of TV, respectively.**

**a,** Lifetime images from traditional WF-fdFLIM, Wiener-SIM reconstruction (raw SIM), Hessian-SIM reconstruction with cross-correlation (CC) regularization, Hessian-SIM reconstruction with total variation (TV) regularization, and Hessian-SIM reconstruction with combined CC + TV regularization (from left to right). All SIM results are reconstructed from DC-combined widefield data.

**b,** Local lifetime standard deviation maps computed over every 4×4 pixels, arranged in the same sequence as in **a**.

**c,** Intensity images corresponding to the five reconstruction methods.

**d,** Normalized intensity line profiles from **c** fitted using a Gaussian function. The spatial resolutions are quantified using the full-width at half-maximum (FWHM).

**e,** Lifetime line profiles extracted from **c** corresponding to the profiles shown in **d**.

**f,** Resolution quantified by decorrelation analysis for the Hessian-SIM reconstruction with combined CC + TV regularization (top), with the corresponding spectra shown on a logarithmic scale (bottom).

**g,** Lifetime histograms and corresponding Gaussian fits for the five reconstruction methods.

Scale bar: 500 nm.

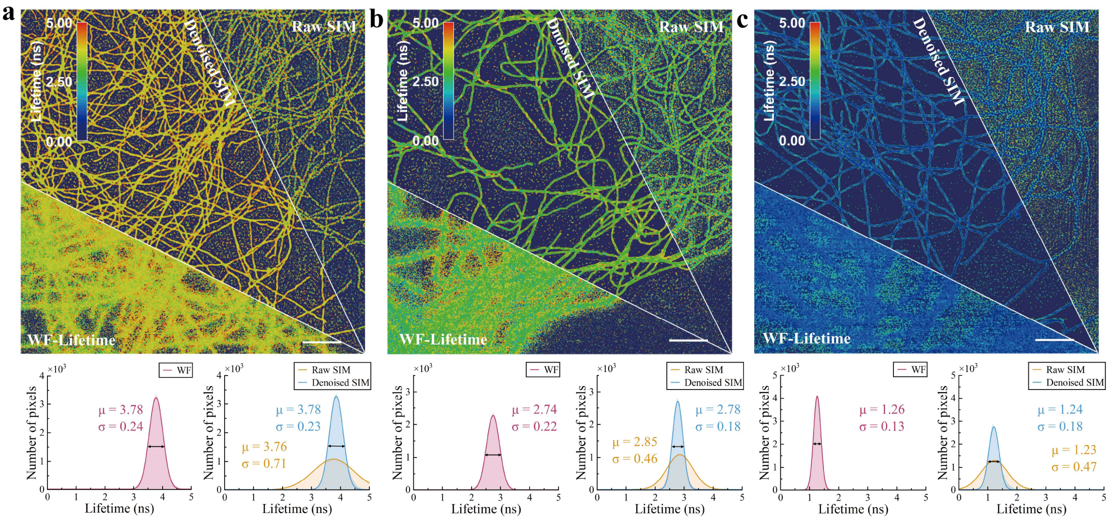


**Supplementary Fig. 7| Lifetime accuracy test of the SIM-fdFLIM system using single-labeled fixed samples.**

**a,** AF488-labeled microtubules excited with a 488 nm laser and detected at ~500 nm.

**b,** AF546-labeled microtubules excited with a 561 nm laser and detected at ~600 nm.

**c,** AF647-labeled microtubules excited with a 638 nm laser and detected at ~700 nm.

Raw SIM denotes SIM results of Wiener reconstruction. Denoised SIM donates Hessian SIM with RIVE. All SIM results are reconstructed from DC-combined widefield data. Red histogram and corresponding Gaussian fits: WF fdFLIM results; Yellow histogram and corresponding Gaussian fits: raw SIM-fdFLIM results; Blue histogram and corresponding Gaussian fits: denoised SIM-fdFLIM results. Scale bar: 2 μm.

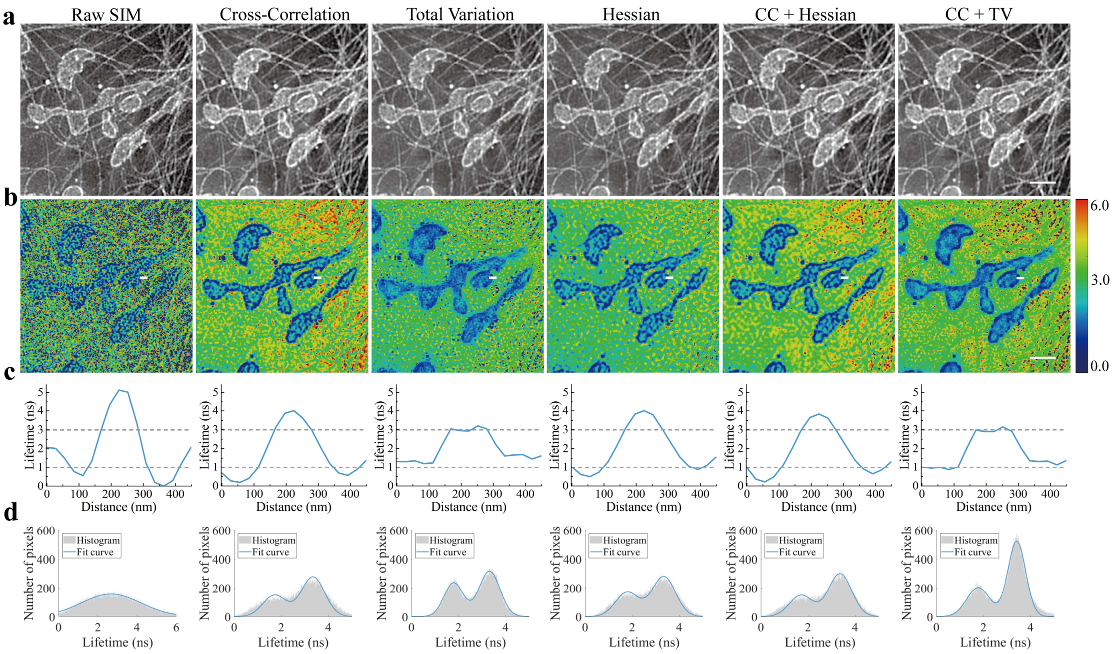


**Supplementary Fig. 8| Effect of denoising regularization on reconstruction results.**

**a,** Reconstructed intensity images obtained by: Wiener-SIM reconstruction only (raw SIM); Hessian-SIM reconstruction with cross-correlation (CC) regularization; Hessian-SIM reconstruction with total variation (TV) regularization; Hessian-SIM reconstruction with Hessian regularization; Hessian-SIM reconstruction with CC + Hessian regularization; and Hessian-SIM reconstruction with CC + TV regularization (from left to right). All SIM results are reconstructed from DC-combined widefield data.

**b,** Corresponding lifetime images processed using the same set of regularizations as in **a.**

**c,** Lifetime line profiles extracted from the locations indicated in **b.**

**d,** Lifetime histograms of the images in **b**, each fitted with a double-Gaussian distribution.

Scale bar: 2 μm.

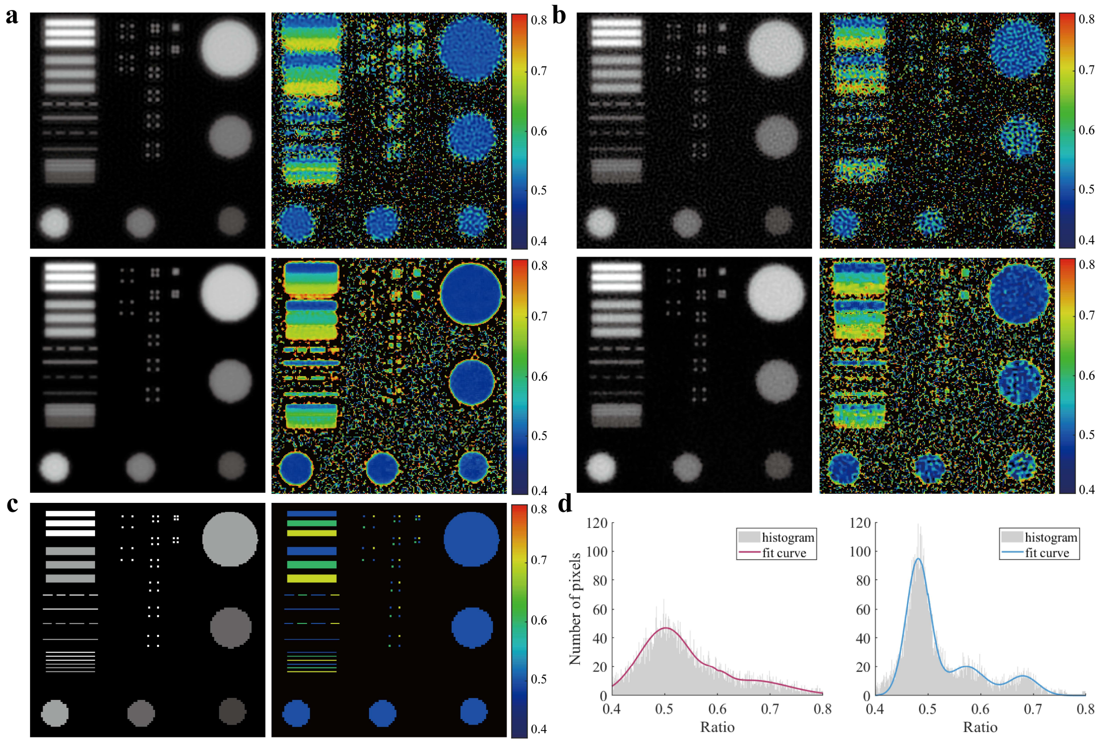


**Supplementary Fig. 9| Simulation illustrating the performance of the denoising algorithm.**

**a-b,** Simulated intensity (left) and ratio (right) reconstructions under Gaussian noise levels of 1% (a) and 5% (b) of the maximum ground-truth intensity. For each condition, raw SIM-fdFLIM (top row) reconstruction and denoised SIM-fdFLIM (bottom row) reconstruction are shown.

**c,** Ground-truth intensity (left) and ratio (right).

**d,** Histograms of ratio images from **b**, comparing raw SIM (left) and denoised SIM (right), each fitted with a three-Gaussian distribution.

**a-c**: Image brightness in the intensity images corresponds to intensity values, with brighter regions indicating higher intensity. Despite varying intensity conditions, the ratio distributions in denoised SIM remain stable.

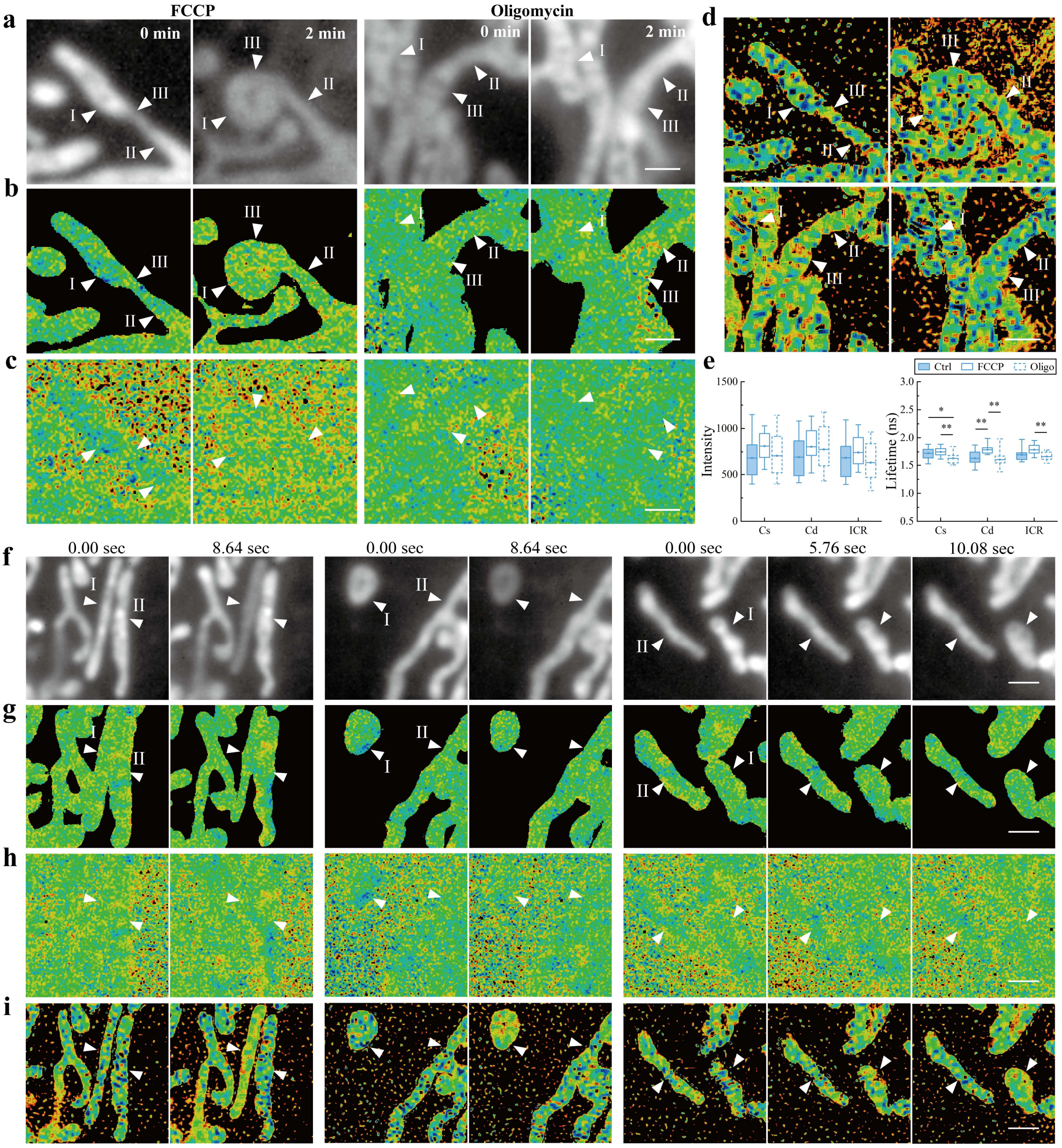


**Supplementary Fig. 10| Various imaging modes of live cells stained with PKMR.**

**a,** WF intensity images of mitochondria treated with FCCP or oligomycin. Arrow I: sparse cristae; II: dense cristae; III: inter cristae region (ICR).

**b,** Widefield lifetime images of mitochondria treated with FCCP or oligomycin, with a mask generated from the intensity image as in **Fig. 3e**.

**c,** Widefield lifetime images of the mitochondria shown in **a**, without mask.

**d,** SIM lifetime images of the mitochondria shown in **a**, without mask.

**e,** Box plots showing the mean intensity (left) and lifetime (right) of mitochondrial internal structures under different conditions. Cs: sparse cristae; Cd: dense cristae; ICR: inter-cristae region. $n \geq 10$ cells.

**f,** Representative mitochondrial WF intensity images exhibiting rapid changes in lifetime and intensity. I: changing mitochondria; II: stable mitochondria. Left: tubular changing mitochondria;

middle: round changing mitochondria; right: tubular-to-round transitioning changing mitochondria.

**g,** WF lifetime image of the mitochondria shown in **f**, with a mask generated from the intensity image as in **Fig 3h**.

**h,** WF lifetime image of the mitochondria shown in **f**, without mask.

**i,** SIM lifetime image of the mitochondria shown in **f**, without mask.

Scale bar: 1 µm. In all lifetime images, background areas with a lifetime > 3 ns were set to black. For all box plots, data are presented as mean (filled circle), median (center line), interquartile range (box), and 5-95% range (whiskers). *P < 0.05; **P < 0.01.

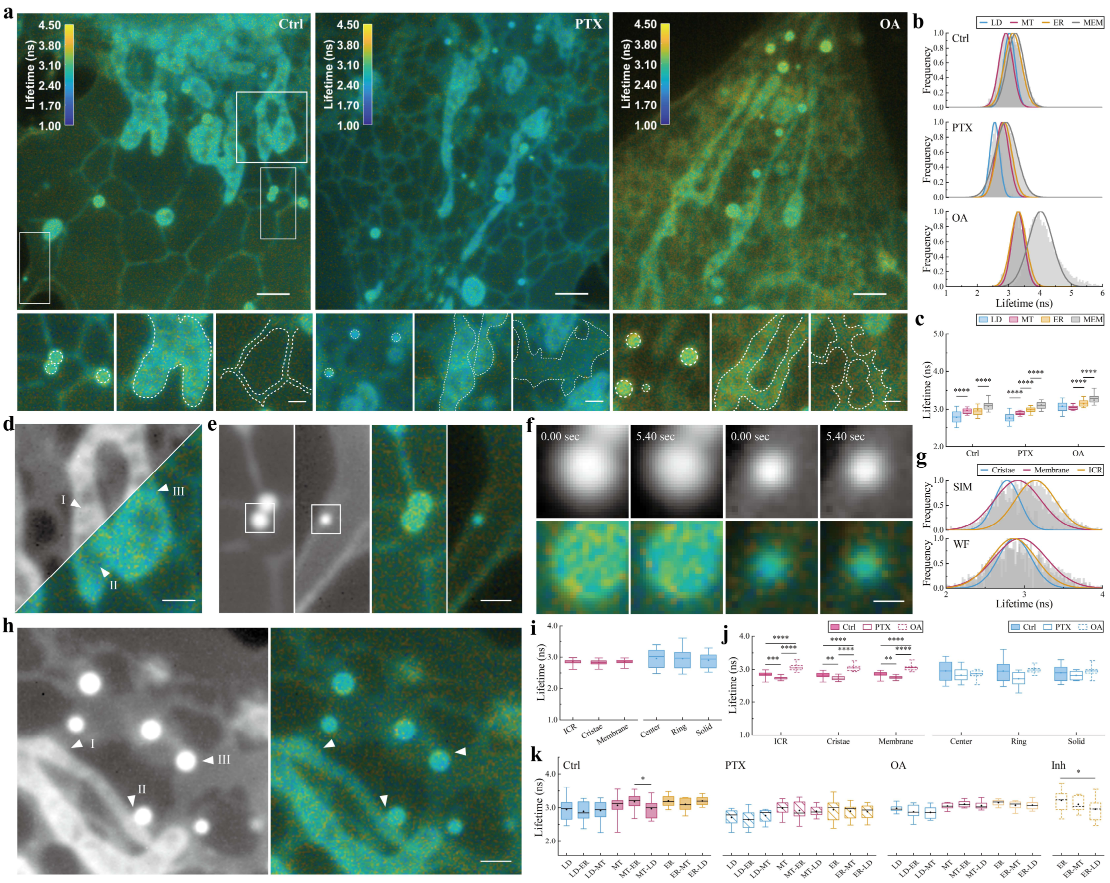


**Supplementary Fig. 11| WF-fdFLIM imaging of live cells stained with Nile Red.**

**a,** Representative intensity-weighted lifetime images of untreated (left, Ctrl), paclitaxel-treated (middle, PTX), and oleic acid-treated (right, OA) cells. The bottom row shows magnified views of lipid droplets, mitochondria and endoplasmic reticulum for each condition. Scale bar: 2 µm; zoom-in panels: 500 nm.

**b,** Lifetime distributions and the corresponding Gaussian fits of lipid droplets (LD), mitochondria (MITO), endoplasmic reticulum (ER) and the plasma membrane-dominant peripheral regions (MEM) under the conditions shown in **a**.

**c,** Box plots showing the distributions of mean lifetimes of LD, MITO, ER, and MEM under different conditions. $n \geq 34$ ROIs.

**d,** Intensity (upper-left) and intensity-weighted lifetime (lower-right) images of mitochondria from the selected regions of interest in **a**. Triangle arrows indicate the mitochondrial cristae (I), ICR (II), and membrane (III) regions. Scale bar: 500 nm.

**e,** Intensity (left) and Intensity-weighted lifetime (right) images of ring-shaped and solid lipid droplets, arranged horizontally. Scale bar: 500 nm.

**f,** Intensity (top row) and intensity-weighted lifetime (bottom row) images of the lipid droplets indicated in **e**, shown at consecutive frames (5.4 s apart). Scale bar: 200 nm.

**g,** Lifetime distributions and the corresponding Gaussian fits of mitochondrial cristae, ICR, and membrane indicated in **d**, acquired by SIM (top) and WF (bottom) imaging.

**h,** Intensity (left) and Intensity-weighted lifetime (right) images of organelle contact regions. Triangle arrows indicate the contact regions between mitochondria and endoplasmic reticulum (I), mitochondria and lipid droplets (II), and endoplasmic reticulum and lipid droplets (III). Scale bar: 500 nm.

**i,** Box plots showing the distributions of mean lifetimes of internal regions in mitochondria (left) and lipid droplets (right). $n \geq 12$ cells.

**j,** Box plots showing the distributions of mean lifetimes of internal regions in mitochondria (left) and lipid droplets (right) under different conditions. $n \geq 12$ cells.

**k,** Box plots showing the distributions of mean lifetimes in 224 nm-diameter contact regions between organelles under different conditions. Inh: DGAT inhibitors treatment group. Labels: A denotes the bulk region of organelle A, whereas A-B denotes the contact region on the A side of the A-B interface. $n \geq 12$ cells.

For all box plots, data are presented as mean (filled circle), median (center line), interquartile range (box), and 5-95% range (whiskers). *P < 0.05; **P < 0.01; ***P < 0.001; ****P < 0.0001.

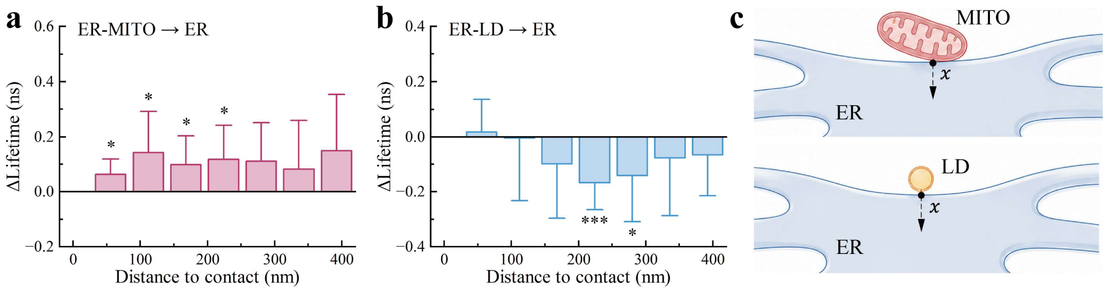


**Supplementary Fig. 12| Distance-dependent lifetime changes around ER-organelle contact sites.**

**a,** Bar plots showing Δlifetime relative to bulk ER as a function of distance from ER-MITO contacts. The x-axis represents the distance between the center of the analyzed 112 nm-diameter region and the ER-MITO contact site. $n \geq 9$ cells.

**b,** Bar plots showing Δlifetime relative to bulk ER as a function of distance from ER-LD contact sites. $n \geq 9$ cells.

**c,** Schematic illustrations of ER-MITO (top) and ER-LD (bottom) contact regions, showing the definition of the measured distance $x$ from the contact site.

For all bar plots, data are presented as mean with one-sided SD error bars. *P < 0.05; ***P < 0.001.

Labels: A-B denotes the contact region on the A side of the A-B interface.

**Supplementary Table 1. Performance comparison of FLIM imaging techniques.**

| Method | Detection strategy | Resolution | Field of view | Imaging rate | Live-cell suitability | Quantitative lifetime preservation/ Main limitation |
|---|---|---|---|---|---|---|
| WF FLIM[1] | Camera-based fdFLIM | Diffraction-limit | Large | Hz-tens of Hz | High | Quantitative but not SR |
| Confocal FLIM[2] | Point-scanning TCSPC | Diffraction-limit (~180-200 nm) | Moderate | Seconds per frame | Medium | Quantitative but scanning-limited |
| STED FLIM[3] | TCSPC + depletion scanning | Nanoscale (< 100 nm) | Small | Minutes per frame | Low | Quantitative but limited by dose and scanning throughput |
| ISM FLIM[4] | SPAD-array image scanning | ~1.4-2× confocal | Moderate | Seconds per frame | Medium | Quantitative but limited throughput |
| SMLM FLIM[5] | TCSPC + Single-molecule localization | Molecular/ localization scale | Very small | Tens of minutes per frame | Low | Quantitative but long acquisition |
| SIM-FLIM (EO)[6] | EO lifetime detection | Lifetime: 156 nm Intensity: 123 nm | 55-μm diameter | 0.1 Hz | Medium | Ratio-domain artefact correction not addressed; custom EO setup |
| SIM-fdFLIM | Camera-based fdFLIM | ~100 nm | 56 × 56 μm² | 2.8 Hz | High | Ratio-preserving SR reconstruction; validated lifetime fidelity |

This table provides a benchmark comparison of representative FLIM modalities. Widefield and confocal FLIM provide quantitative lifetime measurements but remain diffraction-limited or scanning-limited. STED-FLIM, ISM-FLIM and SMLM-FLIM improve spatial resolution but typically sacrifice field of view, acquisition speed or dense live-cell compatibility. SIM-based FLIM partially decouples this trade-off by combining widefield acquisition with SR reconstruction. Among these approaches, SIM-fdFLIM uniquely combines ~100 nm lifetime-map resolution, 2.8 Hz live-cell imaging over a 56 × 56 μm² field of view and ratio-preserving reconstruction that maintains quantitative lifetime fidelity after SR recovery. Values are representative of reported implementations rather than theoretical limits and depend on fluorophore brightness, photon budget, field of view, sample type and reconstruction criteria.


1. Wu, H.-M. *et al.* Widefield frequency domain fluorescence lifetime imaging microscopy (FD-FLIM) for accurate measurement of oxygen gradients within microfluidic devices. *Analyst* **144**, 3494–3504 (2019).
2. Gerritsen, H. C., Asselbergs, M. a. H., Agronskaia, A. V. & Sark, W. G. J. H. M. V. Fluorescence lifetime imaging in scanning microscopes: acquisition speed, photon economy and lifetime resolution. *Journal of Microscopy* **206**, 218–224 (2002).
3. Bianchini, P. *et al.* STED nanoscopy: a glimpse into the future. *Cell Tissue Res* **360**, 143–150 (2015).
4. Castello, M. *et al.* A robust and versatile platform for image scanning microscopy enabling super-

resolution FLIM. *Nat Methods* **16**, 175–178 (2019).
5. Thiele, J. C. *et al.* Confocal Fluorescence-Lifetime Single-Molecule Localization Microscopy. *ACS Nano* **14**, 14190–14200 (2020).
6. Marchand, R. *et al.* Super-resolution structured illumination microscopy of cellular dynamics with fluorescence lifetime contrast. *Optica* **13**, 1065 (2026).

**Supplementary Video 1.** Intensity, lifetime, and lifetime-overlaid intensity images (from top to bottom) of immunolabeled microtubules (AF546) and mitochondria (AF555) in a fixed COS-7 cell, acquired by SIM-fdFLIM. Results are compared with WF imaging, and raw SIM-fdFLIM reconstructed using Wiener filtering (see also **Fig 1e**). Scale bar: 3 μm.

**Supplementary Video 2.** Intensity, lifetime, and lifetime-overlaid intensity images (from top to bottom) of immunolabeled microtubules (AF546) and mitochondria (AF555) in a fixed COS-7 cell, acquired by SIM-fdFLIM. Results are compared with WF imaging and SIM-fdFLIM without DC components. Scale bar: 3 μm.

**Supplementary Video 3.** Time-lapse mitochondrial recording with PKMR. Over ~67 s of imaging, the mean intensity decreases due to photobleaching, whereas the mean lifetime remains stable. This demonstrates the robustness of lifetime readout against photobleaching and low signal-to-noise ratio (SNR). Scale bar: 2 μm.

**Supplementary Video 4.** Fast imaging of PKMR-labeled mitochondria at 2.8 Hz, showing dynamic processes including mitochondrial fission, fusion, and nanotunnel extension. Intensity and lifetime images are displayed, with lifetime images masked based on the corresponding intensity images. Scale bar: 5 μm; zoom-in panels: 1 μm.
